# fMRI Informed Montage Selection for Transcranial Electrical Stimulation: Frontoparietal Synchronization for Drug Cue Reactivity

**DOI:** 10.1101/2021.04.12.439544

**Authors:** Ghazaleh Soleimani, Rayus Kupliki, Jerzy Bodurka, Martin Paulus, Hamed Ekhtiari

## Abstract

**Background:** Frontoparietal network (FPN) with multiple cortical nodes is involved in executive functions. Transcranial electrical stimulation (tES) can potentially modulate interactions between these nodes using frontoparietal synchronization (FPS). Here we used fMRI and computational head models (CHMs) to inform electrode montage and dosage selection in FPS.

**Methods:** Sixty methamphetamine users completed an fMRI drug cue-reactivity task. Two sets of 4×1 HD electrodes with anode over F3 and F4 were simulated and spheres around maximum electric field in each hemisphere were defined as frontal seeds. Using frontal seeds, a task-based functional connectivity analysis was conducted based on a seed-to-whole brain generalized psychophysiological interaction (gPPI). Electrode placement for parietal sites was selected based on gPPI results. Task-based and resting-state connectivity were compared between fMRI-informed and classic F3-P3/F4-P4 montages.

**Results:** Whole-brain gPPI showed two significant clusters (left: 506 voxels P=0.006, right: 455 voxels P=0.016), located in the inferior parietal lobule under the CP5 and CP6 electrode location. Pair-wise ROI-based gPPI comparing informed (F3-CP5/F4-CP6) and classic (F3-P3/F4-P4) montages showed significant increased PPI and resting-state connectivity only in the informed montage. Cue-induced craving score was also correlated with left (F3-CP5) frontoparietal connectivity in the fMRI-informed montage.

**Conclusion:** This study proposes an analytic pipeline to select electrode montage and dosage in dual site tES using CHMs and task-based connectivity. Stimulating F3-F4 can tap into both FPN and saliency network (SN) based on the montage selection. Using CHM and fMRI will be essential to navigating ample parameter space in the stimulation protocols for future tES studies.

**Highlights:**

- We demonstrated a methodology for montage selection in network-based tES
- Task-based functional connectivity can inform dual-site tES montage selection
- Head models can help to induce balance tES dose in targeted brain regions
- Targeting DLPFC with tES can tap into both saliency and frontoparietal networks
- Lower resting-state frontoparietal connectivity before cue exposure followed by a greater craving

## Introduction

Functional imaging of the human brain has shown that connectivity or synchronization between frontal and parietal parts of the brain, which compose the frontoparietal network (FPN), plays a crucial role in goal-driven behavior and cognitive functions [1–4]. It is assumed that the FPN contributes to the top-down control and self-regulation process by allowing people to control their emotions, behaviors, and desires [5–7]. Furthermore, abnormalities in FPN were reported in many diseases, including schizophrenia [8, 9], depression [10, 11], anxiety [12, 13], obsessive-compulsive disorder (OCD) [14], and substance use disorder [15].

In this context, there is evidence to suggest that non-invasive brain stimulation methods such as transcranial electrical stimulation (tES) can synchronize oscillatory activity or modulate functional connectivity between remote cortical regions such as frontal and parietal nodes in FPN [16–18]. Modulating FPN nodes by increasing phase-amplitude coupling showed promising results in improving executive functions and cognitive performance [19–21]. However, there is subtle nuance in the application of tES for modulating brain networks. One aspect that complicates network-based modulation is multi-focal stimulation by tES which requires determining the ideal configuration of electrodes to optimally target distinct network nodes [22].

High-definition tES, where return electrodes are placed close to the active electrode to produce focal electric field (EF), could be used to simultaneously stimulate multiple nodes of a large-scale network [23]. An overarching consideration in optimizing multi-focal tES is dose selection, including (1) electrode montage and (2) current intensity. In order to inform electrode montage, it has been assumed that electrical stimulation prefers to modulate specific forms of ongoing brain activity [24–27]. In this way, at the network level, tES may preferentially modulate brain networks that are already activated (e.g., by a specific task) [28, 29], and only those active pathways would benefit from the induced EFs [24, 30]. Previous findings suggest that functional magnetic resonance imaging (fMRI) allows us to define cortically extended tES targets based on visualizing functional activity within the networks to inform the ideal site for applying network-based stimulation [22, 31]. Furthermore, for current intensity considerations, gyri-precise computational head models (CHMs), that predict how focal brain stimulation will propagate through the networks, suggest that in multi-site tES studies, anatomical features of each target should be considered separately to ensure a satisfactory balance between tES-induced EF strength in stimulation sites [32]. However, for FPN modulation, most of the published studies so far have placed electrodes over F3/F4 (frontal) and P3/P4 (parietal) locations to guide electrical current to the main nodes of FPN without any further considerations about brain function or brain structure [16, 33–38].

In this study, we used CHMs and task-based connectivity to specify a target map on the cortical surface for modulating FPN. Sixty methamphetamine users did an fMRI drug-cue reactivity task after a short period of abstinence. The potential task-based functional connectivity between frontal and parietal regions was investigated by performing seed-to-whole brain generalized psychophysiological interaction (gPPI) analysis. The electrode placement for the frontal site was selected based on maximum EFs, and the electrode location in the parietal site was determined based on the gPPI results. Both task-based and resting-state functional connectivity were compared between the fMRI informed montage and the classic F3/F4-P3/P4 montage which is commonly used in frontoparietal synchronization (FPS). Lastly, we discussed considerations regarding current intensity in each stimulation site and inter-individual variability based on CHMs. Taken together, here, for the first time, we have suggested an analytic pipeline to inform both montage and current intensity to modulate FPN in the context of a cognitive function of interest (drug cue reactivity) to obtain a clinical outcome (reduction in drug craving) using both brain structure and brain function.

## Materials and Methods

### Participants

Participants included 60 subjects (all-male, mean age ± standard deviation (SD) = 35.86 ± 8.47 years ranges from 20 to 55) with methamphetamine use disorder (MUD). All participants were recruited during their early abstinence from the 12&12 residential drug addiction treatment center in Tulsa, Oklahoma. Human research conducted in this study was in accordance with the Declaration of Helsinki, and all methods were carried out under relevant guidelines and regulations. Written informed consent was obtained from all participants before the scans, and the study was approved by the Western IRB (WIRB Protocol #20171742). More details on inclusion/exclusion criteria are reported in supplementary materials (Section S1). Demographic data can be found in Table 1.

**Table 1.** Demographics and substance use profile. All 60 participants are diagnosed with methamphetamine (meth) use disorder.

| Demographic data | Mean (SD) |
| --- | --- |
| Age (years) | 36.8 (8.7) |
| Education (years) | 13.6 (2.6) |
| BMI | 28.0 (5.1) |
| Age of Meth use onset (years old) | 20.7 (7.1) |
| Duration of Meth use at least once a week (years) | 13.2 (19.8) |
| Cost of Meth (dollar per month) | 1039.5 (1267.1) |
| Dose of Meth (gram per day) | 1.61 (1.51) |
| Life time history of Meth injection <sup>b</sup> (n (%)) | 46 (77%) |
| Days of drug use in the last month (before starting abstinence) |  |
| Meth | 22.4 (10.0) |
| Alcohol | 9.4 (11.7) |
| Heroin | 3.9 (9.1) |
| Methadone | 0.7 (3.9) |
| Other opioids | 2.6 (6.1) |
| Barbiturate | 0.1 (0.6) |
| Sedative | 2.2 (6.2) |
| Cocaine | 0.2 (0.9) |
| Cannabis | 7.2 (9.8) |
| Hallucinogens | 0.4 (1.9) |
| Inhalants | 0.1 (0.4) |
| Duration of current abstinence (days) | 61.5 (34.8) |
| Craving Changes on VAS (0-100) from before to after Scanning | 20.8 (25.8) |

### fMRI cue-reactivity task and resting-state fMRI

Task-based fMRI data, pictorial methamphetamine cue exposure, with a pseudo-randomized order in a block-design, with two sets (meth and neutral) of distinct but equivalent pictures validated in another study [39], were collected. Each block consisted of a series of 6 photos of the same category (meth vs. neutral) that were presented for 5 sec each with a 0.2 sec interstimulus interval. In-between each block, a fixation cross was presented for 8 to 12 sec. The whole task incorporated four neutral and four meth picture blocks (i.e., 8 blocks in total with 6 pictures of each category in every block). It took approximately 6 min (The fMRI task codes and stimuli are available at https://github.com/rkuplicki/LIBR_MOCD). Resting-state fMRI data were also collected from all participants before the task-based fMRI and after the structural scans. During resting-state data acquisition, individuals were asked to fix their eyes on the screen (looking at a cross point) and not think about anything in particular. More details on MRI acquisition parameters (Section S2) and fMRI preprocessing steps (Section S3) can be found in supplementary materials.

### Seed definition in the frontal lobe: a computational head model approach

F3/F4 electrode location in the EEG 10-10 standard system is commonly used for stimulating frontal brain areas [40, 41]; however, suggesting an optimum site in the parietal lobe for inducing FPS based on a priori brain mapping knowledge has not been introduced yet. To find an appropriate location in the parietal lobe, in the first step, frontal regions were targeted. Typical high definition (HD) electrode set-ups utilize a 4×1 ring configuration were simulated, where four return electrodes over AF3, F1, F5, and FC3 surrounded the centered electrode over F3 [42, 43]. After electrode placement, EFs were calculated based on the finite element method (FEM) using SimNIBS 3.2 software with a total current strength of 2 mA in the anode and 0.5 mA in each return electrode [44]. Ernie’s head mesh (the individual head model of the normal adult male included in SimNIBS) was used to create the CHM[45]. After the calculation of EFs in the subject space, EFs were transformed into the MNI space. Then, the highest EF location was determined, and a sphere with r = 10 mm was placed around the highest EF location. All of these steps were replicated for the right hemisphere by placing the anode electrode over F4 and cathodes over AF4, F2, F6, and FC4 and a 10 mm sphere was defined around the maximum EF location in the right hemisphere in MNI space. These spheres, which are considered frontal seeds, were combined with the MNI mask to ensure analyses did not include signals from non-brain or white matter voxels.

### Seed definition in parietal lobe: Seed-to-voxel psychophysiological approach

Brain regions whose functional connectivity with the defined frontal masks differs during exposure to meth and neutral cues were determined as parietal targets. CONN as a functional connectivity toolbox was used to perform generalized psychophysiological interaction (gPPI) analysis [46]. The gPPI analysis is a task-based modulation in connectivity that identifies brain regions whose connectivity with a seed region (in this case, frontal masks) varies as a function of psychological context (in this case, presenting either meth or neutral stimulus). Above mentioned bilateral frontal masks obtained from the highest EFs were considered separately as seed regions of interest (ROIs) to examine whether and how meth vs. neutral cues alter the functional coupling between the frontal area and other parts of the brain.

The seed-to-voxel approach was used for gPPI analysis on the meth > neutral contrast. Predictors of the whole-brain-wise gPPI analyses included the time course of the task (psychological term), the time course of a frontal mask as seed region (physiological term), interactions between psychological and physiological terms (PPI term), and covariates [47]. First, an averaged BOLD time course across selected voxels was extracted for each frontal mask and used as a physiological regressor for each individual. For the first-level analysis, a PPI regressor (interaction term) was generated for each condition as the element-by-element product of the ROI time series and coding for the task effect (boxcar function of meth and neutral cues). For frontal seeds, meth > neutral contrast images were entered into a regression model at the second level.

Significant clusters obtained from gPPI analysis were used for defining parietal ROIs. The t-map results and 10-10 EEG electrode positioning system were projected over the cortex in the MNI space. Distance between centers of active clusters and all EEG electrode locations were calculated, and the nearest electrodes to the active clusters were determined. Then, 10 mm spheres were defined around the location of the nearest electrodes over the cortex. These spheres, as parietal masks, were combined with the MNI mask to ensure analyses did not include signals from non-brain or white matter voxels. EF distribution patterns were also simulated using two HD electrode configurations with anode electrodes over parietal masks to show whether tES-induced EF can strongly stimulate defined parietal regions.

### ROI-to-ROI task-based modulation in functional connectivity: ROI-to-ROI gPPI analysis

The ROI-to-ROI gPPI analysis was performed to determine how frontal and parietal regions interact in a task-dependent manner. Frontal and parietal masks in the right and left hemispheres, defined in previous sections, were considered as four separate ROIs. Physiological, psychological, and interaction terms were determined as described in the seed-to-whole brain gPPI analysis. Pair-wise gPPI computation was made for every possible pair-wise combination of selected ROIs for each individual. At the group level, random effect analysis was used across participants, and the one-sample t-test was calculated to compare ROI-based connectivity for meth vs. neutral conditions.

To compare our results with classic FPS protocols, ROI-to-ROI gPPI analyses were also performed with F3, P3, F4, and P4 ROIs. Frontal masks were identical to the previous frontal masks. However, parietal ROIs were defined based on the area under P3 and P4 EEG electrodes. P3 and P4 locations were projected to the cortex in MNI space, and a sphere with r = 10 mm was defined around each site. Non-brain or white matter voxels were removed based on combining these spheres with the MNI mask. ROI-to-ROI gPPI analysis was repeated for the classic ROIs in FPS protocols to determine interactions between these four targets (F3, P3, F4, and P4) during a cue-reactivity task.

### ROI-to-ROI resting-state functional connectivity: ROI-to-ROI correlation analysis

Besides PPI connectivity (task-based regression analysis using gPPI as effective connectivity), ROI-to-ROI resting-state functional connectivity (correlation analysis during rest) was also checked. Left and right seeds in the frontal and parietal masks, for both suggested locations in this study as well as classic electrode placement in FPS protocols, were used for ROI-to-ROI resting-state functional connectivity.

### Individualized computational head models

In order to show whether tES-induced EF can stimulate predefined frontal and parietal masks in a balanced ratio, EF distribution patterns were simulated for all sixty participants based on creating CHMs for each individual with four different electrode placements with the anode over: (1) F3 and P3, (2) F3 and CP5, (3) F4 and P4, and (4) F4 and CP6; and cathode electrodes arranged circularly around each anode; between electrode distance and cathode location were identical across the population. The location of the F3, F4, P3, P4, CP5, and CP6 in EEG 10-10 standard system was mapped over the gray matter (GM) in the subject space, and a 10 mm sphere was defined around each location. Averaged EFs in spheres were calculated for each individual to compare EF strength between stimulation sites. Based on the EFs across the population, some suggestions were also made for each stimulation site’s current intensity to have balanced EFs in the frontal and parietal regions using the informed electrode arrangement. More details on the main steps of generating personalized CHMs and group-level analysis of the tES-induced EFs can be found in supplementary materials (Section S4).

### Exploratory connectivity analysis

An exploratory ROI-to-ROI connectivity analysis was performed targeting alterations in indirect modulatory pathways by using two large-scale networks; frontoparietal (FPN) and ventral attention (VAN) (saliency network) networks in Yeo7 atlas [48]. FPN and VAN were investigated since they have central nodes next to the defined masks in classic and informed montages. Since each network in the Yeo7 atlas covered a widely distributed area, Schaefer-400 atlas [49] was applied for extracting the finer sub-regions of each large-scale network and main network nodes in the frontal and parietal parts of the brain were extracted from these networks. Furthermore, brain areas that contribute in top-down regulation (i.e., sub-cortical areas (amygdala, and basal ganglia (nucleous accumbens, ventral and dorsal caudate, ventromedial and dorsolateral putamen, and globus pallidus)), cingulate gyrus and insular cortex [50, 51]) were also extracted using sub-regions in Brainnetome atlas (see supplementary materials section S4) [52]. Task-based and resting-state functional connectivity were calculated between predefined frontal seeds and these extracted nodes of interest.

### Behavioral data

Cue-induced craving was assessed by measuring the change in craving from before to after cue presentations. Participants reported their drug cravings using the Visual Analogue Scale (VAS) score immediately before and after the MRI session. Pearson correlations between drug craving elicited by a cue-reactivity task and frontoparietal connectivity during task-based and resting-state were calculated in both informed and classic montages.

### Statistical results

All of the statistical results for connectivity analyses (task-based or resting-state) are based on the CONN toolbox. In seed-to-whole brain gPPI analysis, active clusters were reported only when surviving a voxel-level statistical threshold of P uncorrected < 0.001 (two-sided *t*-value > 3.46) and a cluster-level threshold of cluster-size > 60 voxels and clusters with false discovery rate (FDR) corrected P < 0.05 were considered as significant clusters. In ROI-to-ROI gPPI analysis, results were reported when survived P uncorrected < 0.05 (two-sided *t*-value > 2). All P values were corrected for the multiple comparison error with false discovery rate (FDR) correction, and both corrected and uncorrected P values were reported in ROI-to-ROI tables. The P values were not corrected in exploratory findings and behavioral data analysis, and only uncorrected P < 0.05 values were reported to show overall trends.

## Results

### Seed definition in the frontal lobe

Ernie’s head mesh was used to create the CHM, and the highest EFs were calculated to determine frontal seeds. As shown in Figure 1, after creating the head model and transformation to the MNI space, a sphere with a 10 mm radius was defined around each maximum value and combined with an MNI mask to remove non-brain and white matter voxels. These spheres are shown with green spheres in Fig.1. Center coordinate (x, y, z in MNI space) for the left frontal side (under F3 electrode) was (−41, 47, 27) and for the right frontal side (under F4 electrode) was (43, 47, 25).

**Figure 1:**
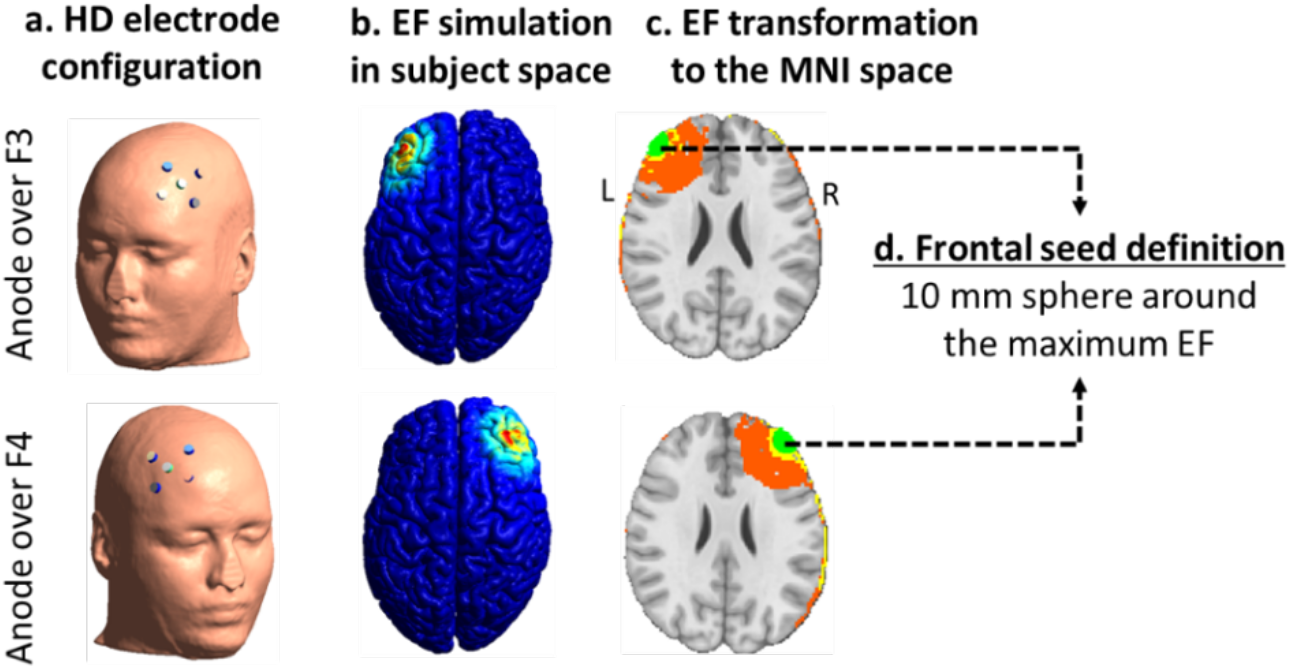
Seed definition in the frontal lobe. **(a).** HD electrode configuration was placed over the frontal lobe in the left (first line) and right (second line) hemispheres separately, with four cathode electrodes circularly placed around each anode (electrode location for the left hemisphere: anode over F3, cathodes over AF3, F1, F5, and FC3. In right hemisphere: anode over F4, cathodes over AF4, F2, F6, and FC4). Current densities were 2 mA for the anode and 0.5 mA for each of the four cathode electrodes. **(b).** Individualized computational head models were generated for each electrode configuration. **(c).** Simulated electric fields (EFs) were transformed into the MNI space. **(d).** The maximum EF was determined, and a sphere with r = 10 mm was placed around the highest EFs in each hemisphere. These spheres were combined with the MNI mask to delete non-brain regions or white matter parts of the spheres. Final masks in the frontal lobe are depicted with green spheres. Abbreviation: EF = electric field; R: right side, L: left side, and left = real left side of the brain.

### Seed selection in the parietal lobe

Seed to whole-brain gPPI analysis was performed using frontal seeds (Fig.2). After setting a threshold (t-value > 3.1 and cluster-size > 60), our results showed that when the frontal seed in the left hemisphere was used, gPPI analysis revealed a cluster survived FDR correction, which is located in the left parietal lobe (x,y,z center in MNI: (−54, −50, 26)). Accordingly, when the frontal seed in the right hemisphere was used, gPPI analysis revealed a cluster survived FDR correction in the right hemisphere, which is located in the parietal lobe (x,y,z center in MNI: (50, −50, 22)). Some other clusters were also found in both gPPI analyses using the left and right seeds (Table 2). However, other clusters did not survive FDR correction.

**Figure 2:**
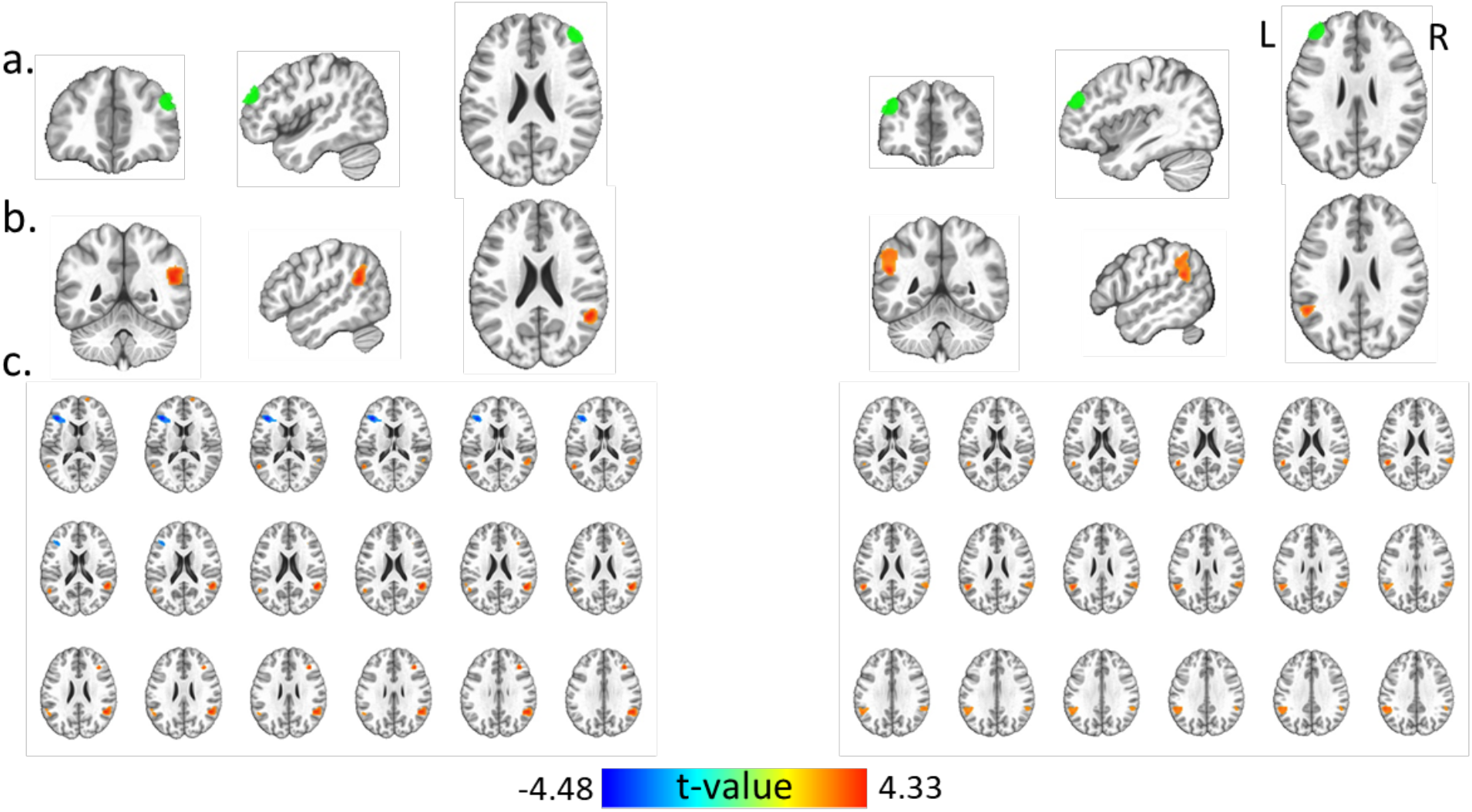
Frontal seed-to-whole brain gPPI results. **(a).** Frontal seeds in the left and right hemispheres are depicted with green spheres; right frontal seed center in MNI space: (43, 47, 25) and left frontal seed center in the MNI space: (−41, 47, 27). Each frontal seed was used for seed-to-whole brain gPPI analysis. **(b).** Frontal seed in each hemisphere showed a significant cluster survived FDR correction (*P* FDR corrected < 0.05, t-value > 3.1, two-sided, cluster size > 60) in the parietal lobe in the same hemisphere; left hemisphere: size = 506 with center (−54, −50, 26) in MNI space (located in IPL caudal area), significant cluster in the right hemisphere: size = 455 with center (50, −50, 22) in MNI space (located in IPL rostroventral area). **(c).** All of active clusters (significant and non-significant) based on gPPI analysis with (t-value > 3.1, two-sided, cluster size > 60). <u>Abbreviation</u>: R: right side, L: left side, and left = real left side of the brain.

**Table 2:** Active clusters in seed-to-whole brain gPPI analysis (meth>neutral) based on using F3/F4 seeds. Activations with P uncorrected < 0.001 at the voxel-level and cluster-size > 60 at the cluster-level are reported in the table. Abbreviation: IPL: inferior parietal lobule, IFG: inferior frontal gyrus, IFS: inferior frontal sulcus, MTG: middle temporal gyrus, STS: superior temporal sulcus, STG: superior temporal gyrus, INS: insular gyrus, MFG: middle frontal gyrus, INS (vId/vIg): ventral dysgranular and granular parts of the insula.

| Seed coordinate<br>(x, y, z) | hemisphere | Location in<br>Brainnetome atlas | MNI peak<br>coordinate |  |  | Cluster<br>Size | t-value<br>in<br>peak | Cluster-level<br>uncorrected<br>*P*-value | Cluster-level<br>FDR<br>corrected<br>*P*-value |
| --- | --- | --- | --- | --- | --- | --- | --- | --- | --- |
|  |  |  | x | y | z |  |  |  |  |
| **F3**<br>(-41, 47, 27) | L | IPL (caudal) | -54 | -50 | 26 | 506 | 4.15 | 0.0003 | **0.006\*\*** |
|  | R | IPL (caudal) | 60 | -46 | 28 | 227 | 3.62 | 0.0079 | 0.091 |
|  | L | Cerebellum | -14 | -92 | -22 | 123 | 4.45 | 0.0402 | 0.308 |
| **F4**<br>(43, 47, 25) | R | IPL (rostroventral) | 50 | -50 | 22 | 455 | 4.30 | 0.0006 | **0.015\*** |
|  | L | IFG (IFS) | -40 | 30 | 16 | 246 | -4.48 | 0.0055 | 0.077 |
|  | R | MTG (anterior STS) | 46 | -32 | -6 | 163 | 3.89 | 0.0230 | 0.214 |
|  | L | pSTS (posterior STS) | -62 | -40 | -2 | 108 | 3.56 | 0.0569 | 0.330 |
|  | L | MTG (dorsolateral) | -56 | -58 | 16 | 106 | 3.93 | 0.0590 | 0.330 |
|  | L | STG (caudal) | -44 | -30 | -4 | 95 | 3.99 | 0.0719 | 0.335 |
|  | R | INS (vld/vlg) | 38 | 32 | 28 | 80 | 4.21 | 0.0956 | 0.356 |
|  | R | MFG (lateral) | 18 | 66 | 10 | 73 | 4.33 | 0.1099 | 0.356 |
|  | L | MTG (aSTS) | -50 | -44 | -18 | 71 | -4.18 | 0.3560 | 0.114 |

\*: $P$ FDR corrected $< 0.05$ ; \*\*: $P$ FDR corrected $< 0.01$

Frontal seeds and significant parietal clusters (located in the left and right IPL) were mapped over the MNI cortex (Figure 3). Calculation distance between the center of active clusters and EEG electrodes showed that CP5 and CP6 are the nearest electrodes to the activations center. Based on the projection of electrode locations from the scalp coordinate to the (x, y, z) coordinate over the cortex in MNI space, CP5 location over the cortex (x, y, z in MNI space) was (−63.94, −46.69, 25.01) and CP6 location was (67.82, −43.78, 26.14). In order to define parietal seeds, 10 mm spheres were placed around CP5 and CP6 coordinates over the cortex in MNI space. The final frontal and parietal seeds are shown in Fig.3.

**Figure 3:**
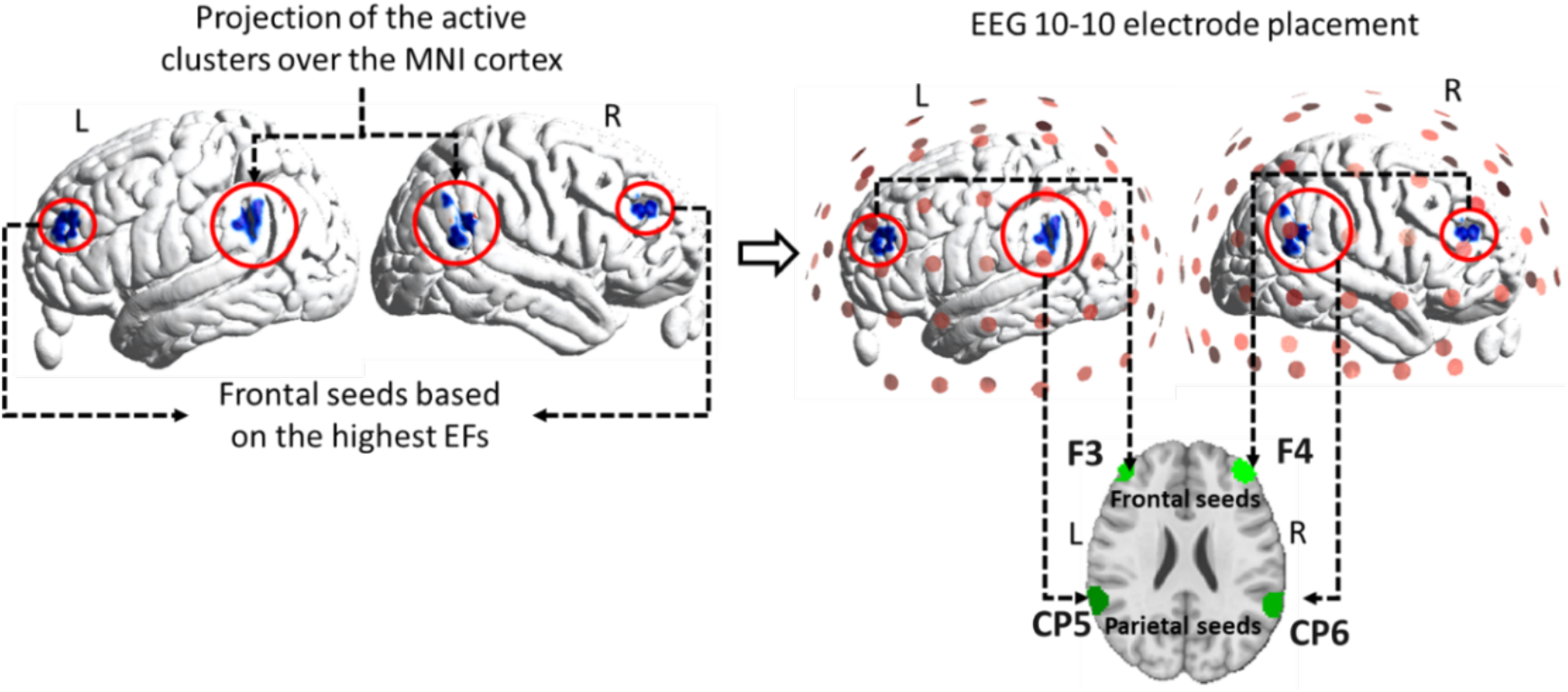
Parietal seed definition. Left panel: projection of the frontal seeds (10 mm spheres around maximum EFs induced by HD electrode configuration over F3 and F4) and active clusters (obtained from frontal seeds-to-whole brain gPPI analysis including IPL caudal area in the left hemisphere and IPL rostroventral area in the right hemisphere) over the cortex in MNI space. Right panel: projection of the EEG 10-10 standard system over the cortex. F3-F4, and CP5-CP6 are the nearest electrodes to the center of frontal masks and active clusters. Parietal seeds were defined based on the 10 mm sphere around the CP5 and CP6 in MNI space. Frontal and parietal seeds are depicted with green spheres. <u>Abbreviation</u>: R: right side, L: left side and left = real left side of the brain.

### ROI-to-ROI task-based modulation in functional connectivity

Based on the frontal and parietal seeds, ROI-to-ROI gPPI analysis was performed to determine the interaction between frontal and parietal ROIs for both informed (F3-CP5/F4-CP6) and classic (F3-P3/F4-P4) montages. Results (meth > neutral) for P uncorrected < 0.05 can be found in Table 2 for task-based functional connectivity and three clusters survived FDR correction in the informed montage. After FDR correction (*P* FDR corrected < 0.05) our results showed increased functional connectivity during the cue-reactivity task between left frontal and right parietal (t value = 3.61), right frontal and right parietal (t value = 2.83), and right frontal and left parietal (t value = 2.68) regions based on the seeds defined for the informed montage. Conversely, in the classic montage, ROI-to-ROI task-based connectivity did not survive FDR correction (*P* FDR corrected > 0.05, Table 3). Connectogram to visualize task-based connections between ROIs in informed and classic montages are shown in Fig.4.

**Table 3:**
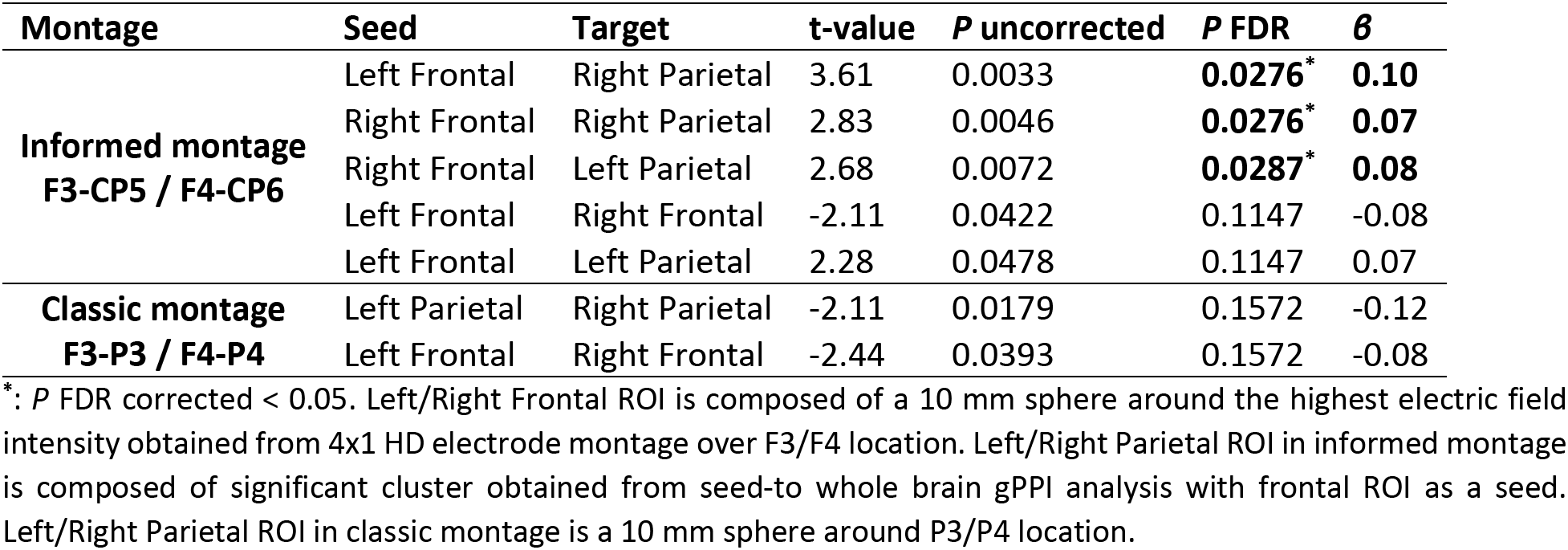
ROI-to-ROI gPPI analysis (meth>neutral) based on using informed (F3-CP5/F4-CP6) and classic (F3-P3/F4-P4) montages. Connections with P uncorrected < 0.05 and averaged beta values for meth vs. neutral conditions are reported in the table.

| Montage | Seed | Target | t-value | $P$ uncorrected | $P$ FDR | $\beta$ |
| --- | --- | --- | --- | --- | --- | --- |
| **Informed montage**<br>**F3-CP5 / F4-CP6** | Left Frontal | Right Parietal | 3.61 | 0.0033 | **0.0276\*** | **0.10** |
|  | Right Frontal | Right Parietal | 2.83 | 0.0046 | **0.0276\*** | **0.07** |
|  | Right Frontal | Left Parietal | 2.68 | 0.0072 | **0.0287\*** | **0.08** |
|  | Left Frontal | Right Frontal | -2.11 | 0.0422 | 0.1147 | -0.08 |
|  | Left Frontal | Left Parietal | 2.28 | 0.0478 | 0.1147 | 0.07 |
| **Classic montage**<br>**F3-P3 / F4-P4** | Left Parietal | Right Parietal | -2.11 | 0.0179 | 0.1572 | -0.12 |
|  | Left Frontal | Right Frontal | -2.44 | 0.0393 | 0.1572 | -0.08 |

\*: $P$ FDR corrected $< 0.05$ . Left/Right Frontal ROI is composed of a 10 mm sphere around the highest electric field intensity obtained from 4x1 HD electrode montage over F3/F4 location. Left/Right Parietal ROI in informed montage is composed of significant cluster obtained from seed-to whole brain gPPI analysis with frontal ROI as a seed. Left/Right Parietal ROI in classic montage is a 10 mm sphere around P3/P4 location.

**Figure 4:**
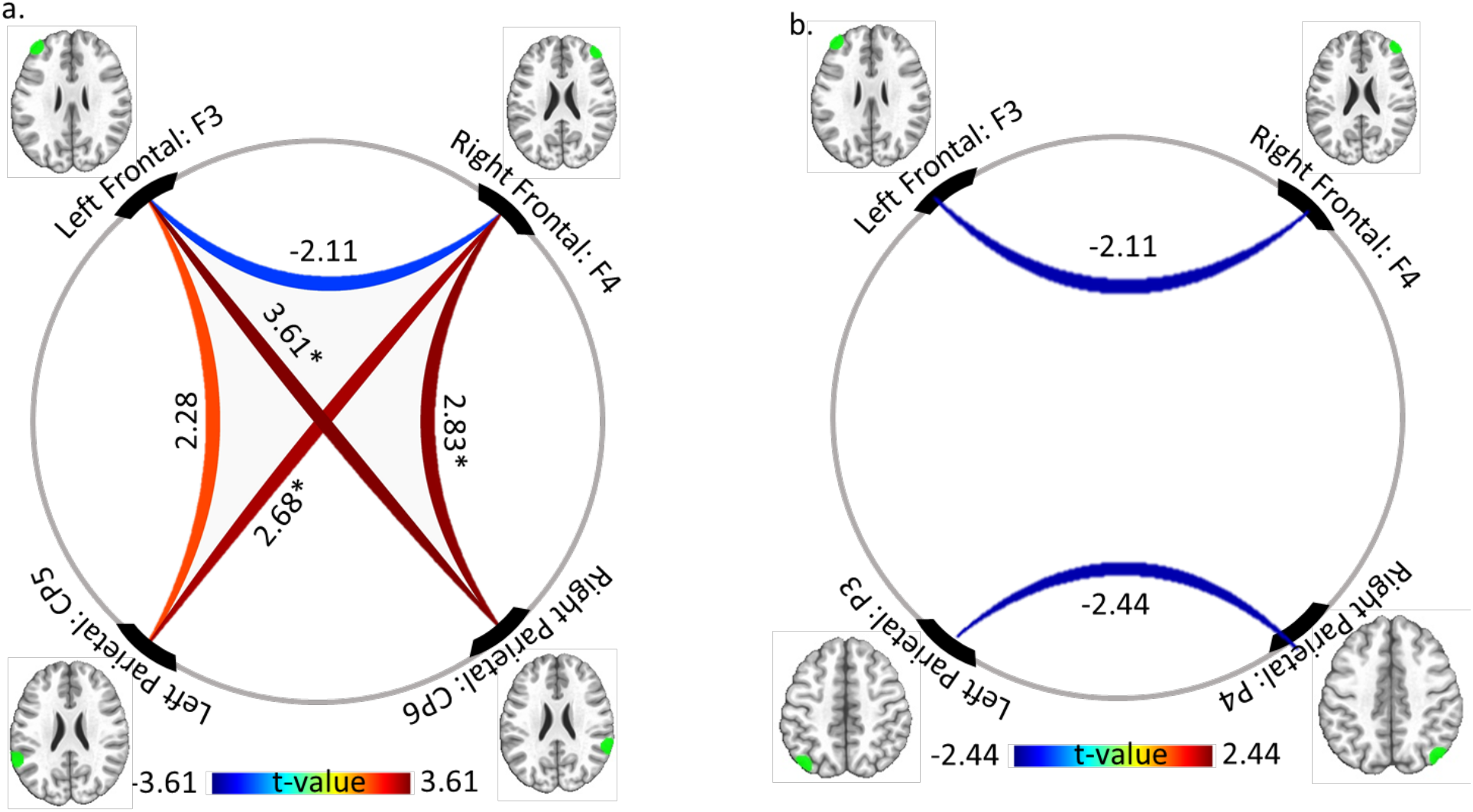
ROI-to-ROI gPPI (meth > neutral) changes using a. F3-CP5/F4-CP6 and b. F3-P3/F4-P4 seeds. Frontal and parietal masks are depicted with green spheres inside the brain based on using F3/F4 masks in the frontal area and CP5/CP6 or P3/P4 masks in parietal regions. Results for PPI connectivity during a cue-reactivity task for meth > neutral contrast are depicted for all connections with P uncorrected < 0.05, and t-values are reported over the pair-wise connections. Colors determine the strength of the connection. Hot colors (yellow and red): increased connestivity and cold color (dark and light blue): decreased connectivity. PPI connectivity survived FDR correction (P FDR corrected < 0.05) are determined with asterisks (‘*’: P FDR corrected < 0.05; ‘**’: FDR corrected < 0.01). Left/Right Frontal ROI is composed of a 10 mm sphere around the highest electric field intensity obtained from 4×1 HD electrode montage over F3/F4 location. Left/Right Parietal ROI in the informed montage is composed of a significant cluster obtained from seed-to whole-brain gPPI analysis with frontal ROI as a seed. Left/Right Parietal ROI in the classic montage is a 10 mm sphere around P3/P4 location.

### ROI-to-ROI resting-state functional connectivity results

ROI-to-ROI resting-state functional connectivity for both informed (F3-CP5/F4-CP6) and classic (F3-P3/F4-P4) montages can be found in Table 4. In the informed montage, connectivity between each pair of the masks was significant (P uncorrected < 0.05) and positive and all of them survived FDR correction (P FDR correction < 0.05).

**Table 4:** ROI-to-ROI resting-state functional connectivity analysis based on using informed (F3-CP5/F4-CP6) and classic (F3-P3/F4-P4) seeds. Connections with P uncorrected < 0.05 and averaged beta values during rest are reported in the table.

| Montage | ROI | ROI | t-value | $P$ uncorrected | $P$ FDR | $\beta$ |
| --- | --- | --- | --- | --- | --- | --- |
| **Informed montage**<br>**F3-CP5 / F4-CP6** | Left Parietal | Right Parietal | 16.14 | 0.0000 | **0.0000** *** | **0.59** |
|  | Right Frontal | Left Frontal | 15.02 | 0.0000 | **0.0000** *** | **0.49** |
|  | Right Frontal | Right Parietal | 4.98 | 0.0000 | **0.0000** *** | **0.16** |
|  | Right Parietal | Left Frontal | 4.16 | 0.0001 | **0.0002** ** | **0.13** |
|  | Left Parietal | Left Frontal | 4.15 | 0.0002 | **0.0003** ** | **0.12** |
|  | Right Frontal | Left Parietal | 2.35 | 0.0223 | **0.0223** * | **0.06** |
| **Classic montage**<br>**F3-P3 / F4-P4** | Right Parietal | Left Parietal | 16.01 | 0.0000 | **0.0000** *** | **0.60** |
|  | Right Frontal | Left Frontal | 15.02 | 0.0000 | **0.0000** *** | **0.49** |
|  | Left Frontal | Left Parietal | -3.94 | 0.0002 | **0.0004** ** | **-0.11** |
|  | Left Frontal | Right parietal | -3.45 | 0.0010 | **0.0016** ** | **-0.09** |

\*: $P$ FDR corrected < 0.05; \*\*: FDR corrected < 0.01; \*\*\*: $P$ FDR corrected < 0.001. Left/Right Frontal ROI is composed of a 10 mm sphere around the highest electric field intensity obtained from 4x1 HD electrode montage over F3/F4 location. Left/Right Parietal ROI in informed montage is composed of significant cluster obtained from seed-to whole brain gPPI analysis with frontal ROI as a seed. Left/Right Parietal ROI in classic montage is a 10 mm sphere around P3/P4 location.

As shown in the lower part of Table 4, our results for classic montage were different from the informed masks in terms of functional connectivity strength and direction of the relationships (positive or negative correlation). In ROI-to-ROI analysis based on F3, F4, P3, and P4 locations in the classic montage,we found significant positive correlations in frontal (between F3-F4) and parietal (between P3-P4) regions (P FDR corrected < 0.05). However, in frontoparietal resting-state connectivity, our results showed negative correlations in F3-P3 and F3-P4 (P FDR corrected < 0.05) ROIs. No significant resting-state functional connectivity was found between right frontal and parietal nodes in classic montage (P FDR corrected > 0.05). Connectogram to visualize resting-state functional connectivity between ROIs in informed and classic ROIs are shown in Fig.5.

**Figure 5:**
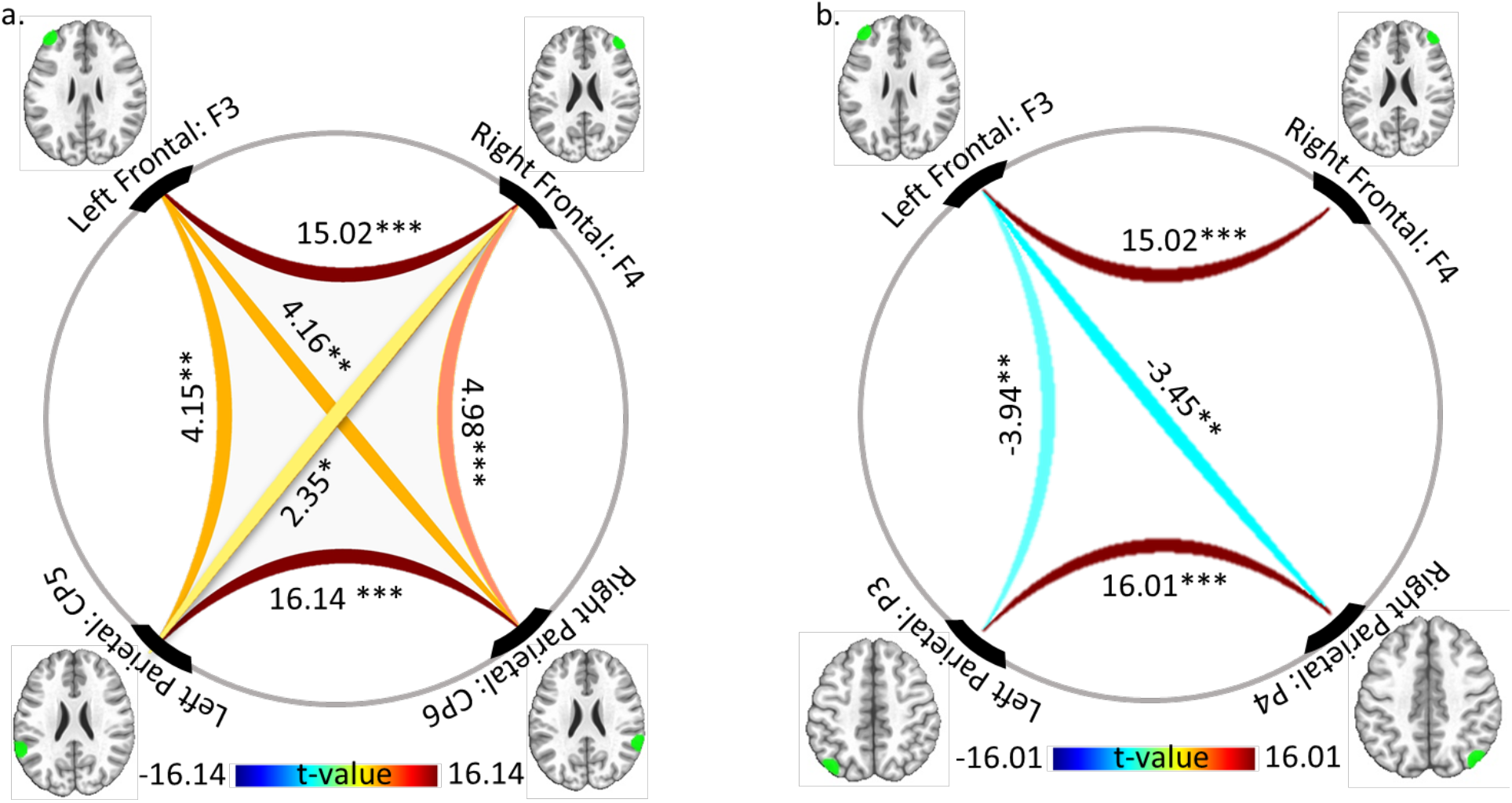
ROI-to-ROI resting-state functional connectivity changes using a. F3-CP5/F4-CP6 and b. F3-P3/F4-P4 seeds. Frontal and parietal masks are depicted with green spheres inside the brain based on using F3/F4 masks in the frontal area and CP5/CP6 or P3/P4 masks in parietal regions. Changes in resting-state functional connectivity are depicted for all connections with P uncorrected < 0.05, and t-values are reported over the pair-wise connection. The connections’ direction and strength are shown using a color bar; hot colors (yellow and red): increased functional connectivity and cold color (dark and light blue): decreased connectivity. Connections survived FDR correction (P FDR corrected < 0.05) are determined with asterisks (‘**\*’**: *P* FDR corrected < 0.05; ‘**\*\*’**: FDR corrected < 0.01; ‘**\*\*\*’**: *P* FDR corrected < 0.001). Left/Right Frontal ROI comprises a 10 mm sphere around the highest electric field intensity obtained from 4×1 HD electrode montage over F3/F4 location. Left/Right Parietal ROI in the informed montage is composed of a significant cluster obtained from seed-to whole-brain gPPI analysis with frontal ROI as a seed. Left/Right parietal ROI in the classic montage is a 10 mm sphere around P3/P4 location.

### Exploratory connectivity results

Exploratory analysis was performed to find task-based and resting-state connectivity between frontal seeds and (1) parietal nodes in two relevant large-scale networks including FPN and VAN based on atlas-based parcellation of the brain regions, as well as (2) other cortical and sub-cortical areas including the insula, cingulate gyrus, basal ganglia, and amygdala that might be modulated through the top-down regulation. In network-based parcellation, as shown in Fig.6 using Yeo7 atlas, frontal and parietal seeds in the classic montage were located near the frontal and parietal nodes in FPN. However, VAN also had frontoparietal nodes, and parietal seeds in the informed montage were located near the main nodes in VAN (instead of FPN). Our results showed that in task-based and resting-state connectivity, frontal seeds had more positive connections with those parietal nodes in VAN and FPN which were located next to the parietal nodes in the informed montage.

**Figure 6:**
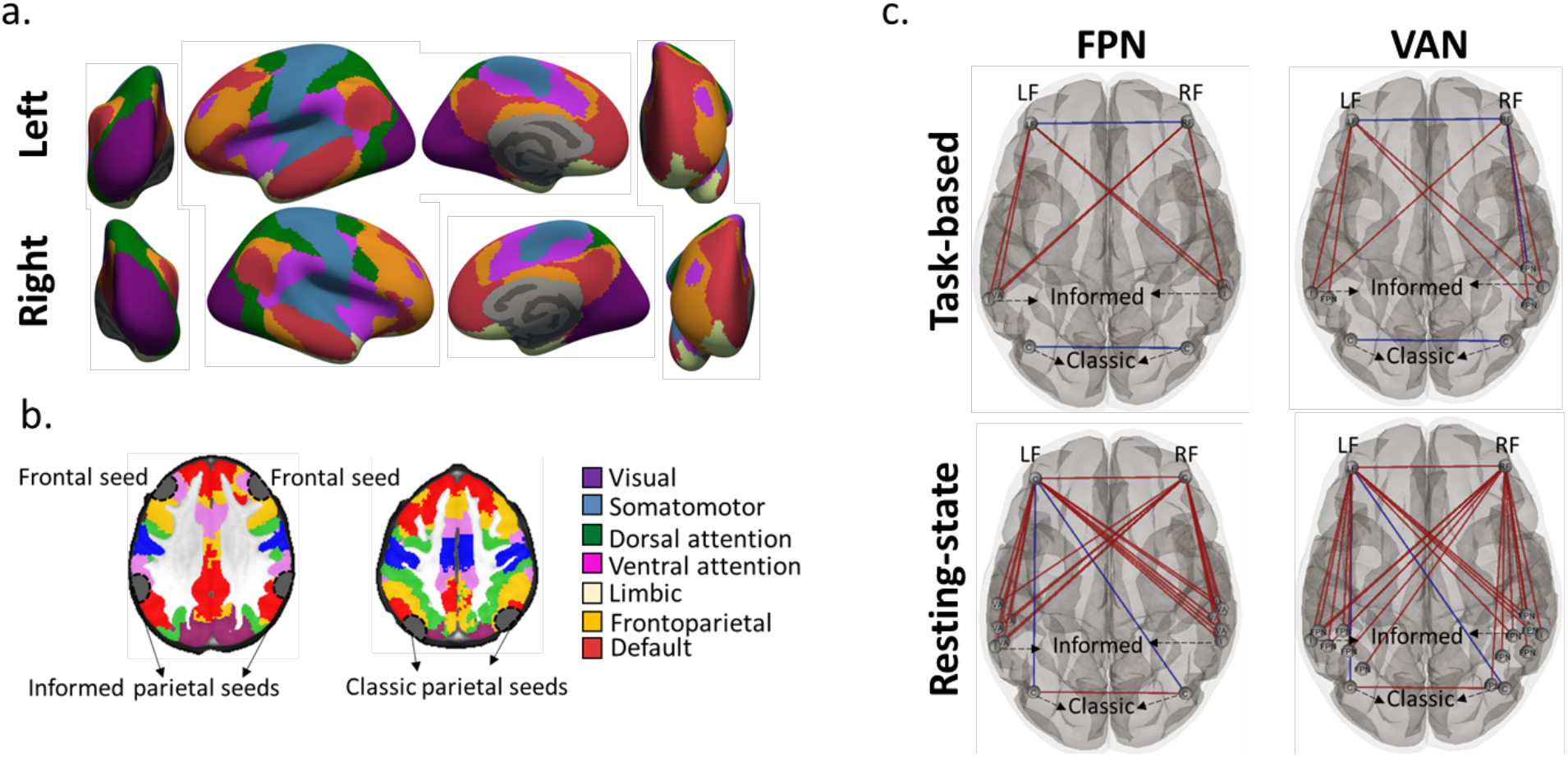
Network-based connectivity: **a.** Large-scale network parcellation is depicted based on Yeo7 standard atlas. **b.** Location of the frontal seeds and parietal seeds in the informed and classic montage with regard to the FPN and VAN. **c.** PPI connectivity (first row) for meth > neutral contrast and resting-state (second row) connectivity between frontal seeds (F3 and F4) and parietal nodes in FPN and VAN are calculated. Parietal nodes in informed and classic montage are also added to the parietal nodes in FPN or VAN. Connection threshold was set to P uncorrected < 0.05. Within network PPI connectivity did not survive correction using FDR method. However, all of resting-state connectivity survived FDR multiple comparisons correction. Red line: increased connectivity and blue line: decreased connectivity. Abbreviation: LF: left frontal; RF: right frontal.

As shown in Fig.7, during a cue-reactivity task, frontal seeds (F3 and F4) revealed interactions with the ventral insula, right nucleus accumbens, ventral caudate, medial amygdala, and caudodorsal part of the anterior cingulate gyrus. During resting-state, functional connectivity changes were found between frontal sites and different parts of the insula, ventral striatum (including ventral caudate and nucleous accumbens), amygdala, and cingulate gyrus.

**Figure 7:**
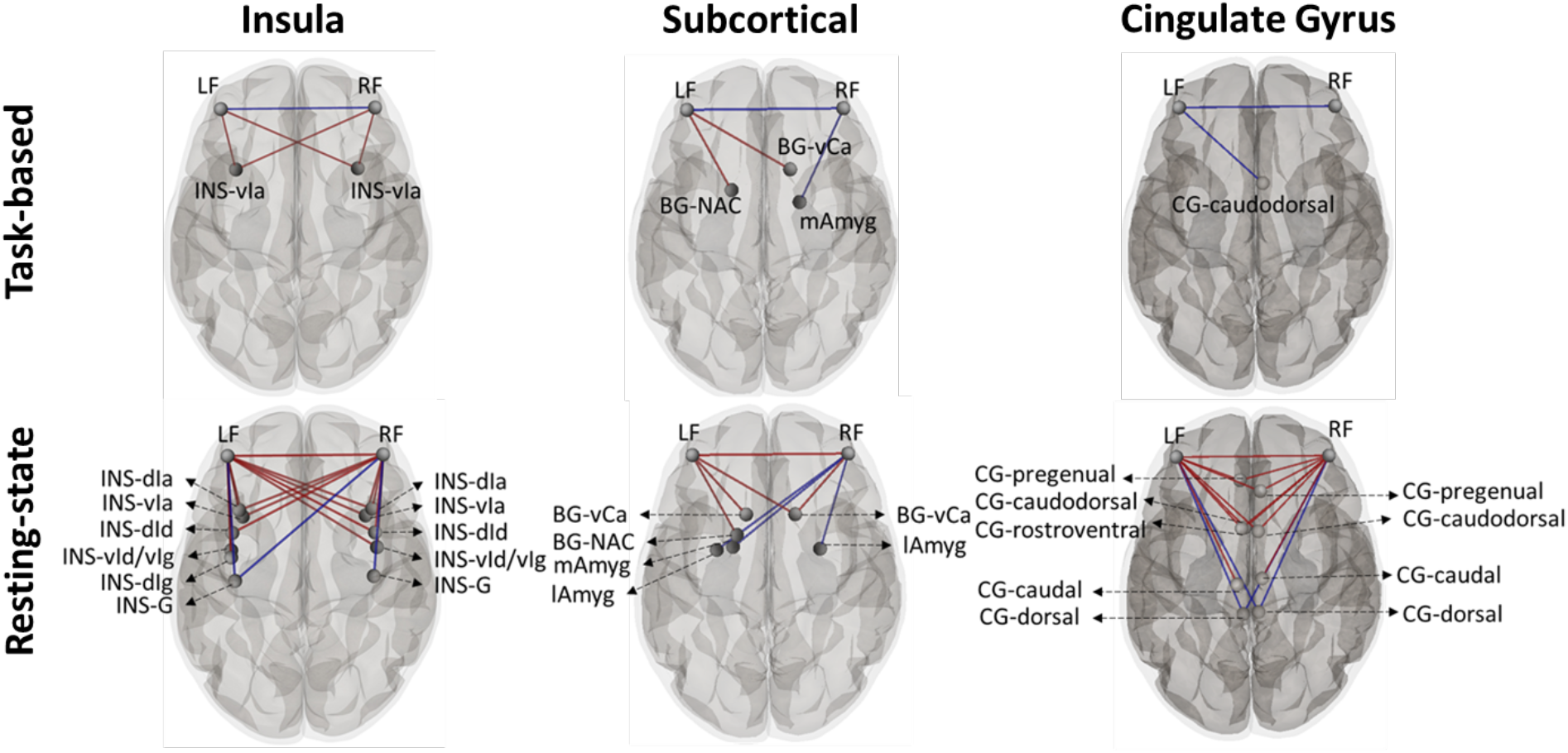
Frontal to cortical-subcortical connectivity: PPI connectivity (first row for meth > neutral contrast and resting-state (second row) functional connectivity between frontal seeds (F3 and F4) and insular sub-regions (first column), basal ganglia (nucleous accumbens, ventral and dorsal caudate, ventromedial and dorsolateral putamen, and globus pallidus) and amygdala second column), and cingulate gyrus (third column) are calculated. Brain parcellation was performed based on the Brainnetome atlas. Connection threshold was set to P uncorrected < 0.05. PPI connectivity did not survive correction using the FDR method. However, all of resting-state connectivity survived FDR multiple comparisons correction. Red line: increased connectivity and blue line: decreased connectivity. Abbreviation: LF: left frontal; RF: right frontal; INS-vIa: ventral angular insula; INS-dIa: dorsal angular insula; INS-dId: dorsal dysgranular insula; INS-vId/vIg: ventral dysgranular and granular insula; INS-dIg: dorsal granular insula; INS-G: hyper granular insula; BG-vCa: basal ganglia ventral caudate; BG-NAA: basal ganglia nucleus accumbens; mAmyg: medial amygdala; lAmyg: lateral amygdala; CG: cingulate gyrus.

All exploratory ROI-wise analysis connection threshold was set to P uncorrected < 0.05 to show the changes’ trends. Task-based connectivity did not survive correction using the FDR method. Conversely, all of the resting-state connectivity, in Fig.6 and Fig.7, survived FDR multiple comparisons correction (P corrected < 0.05).

### Behavioral data analysis results

Craving score based on VAS was significantly (P < 0.001) increased from before (Mean ± SD = 40.95 ± 30.89) to after (Mean ± SD = 61.73 ± 31.11) scanning. As an exploratory finding, as shown by scatterplots in Fig.8, by considering P uncorrected < 0.05, frontal-to-parietal connectivity in the left hemisphere was correlated with craving changes on VAS (0-100) from before to after scanning in the informed montage (F3-CP5) both in task-based connectivity based on meth vs. neutral contrast (r = 0.25, P uncorrected = 0.043) and resting-state (r = −0.28, P uncorrected = 0.031) functional connectivity. Furthermore, a significant negative correlation was found between resting-state and task-based connectivity in the left hemisphere regarding the informed montage (r = −0.31, P uncorrected = 0.014). However, no correlation with P uncorrected < 0.05 was found between cue-induced changes and connectivity within the classic montage.

**Figure 8:**
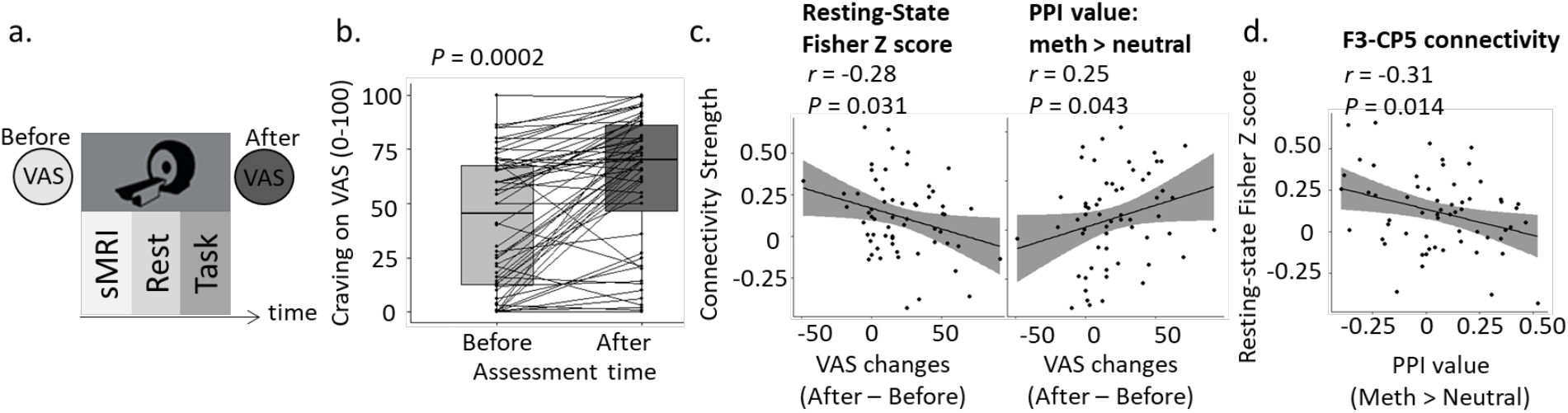
Cue-induced craving changes and frontoparietal connectivity. **a.** Data collection sequence, **b.** VAS reported by all 60 participants before (light gray) and after (dark gray) scanning. Boxplots showing effects of time (before and after drug cue reactivity task) on drug craving and dots represent the data for each individual. P value for differences between VAS before and after scanning is reported above boxplots **c.** Exploratory results for P uncorrected < 0.05 are reported. Dots represent each individual and the grayscale is an indicator of 95% confidence intervals. R and P values are reported for Pearson correlations between craving changes on VAS (0-100) from before to after scanning and resting-state (left side of the figure)/task-modulated (right side of the figure) connectivity in the left hemisphere; informed montage (F3-CP5). Resting-state connectivity refers to correlation between F3 and CP5 masks and PPI value (task-based connectivity) referes to connectivity between F3 and CP5 masks that differs during exposure to meth versus neutral cues. **d.** Scatter plot for pearson correlation between resting-state and PPI values (task-based connectivity) in F3-CP5. Abbreviation: VAS: Visual Analogue Score.

### Group-level analysis of computational head models results

Four different frontoparietal HD montages were simulated for each individual with anodes over (1) F3 and P3, (2) F3 and CP5, (3) F4 and P4, and (4) F4 and CP6. Results of two first montages with anodes over the left hemisphere are represented in Fig.9. As shown in Fig.9, when 2 mA current intensity in the anode and 0.5 mA in each cathode were used for stimulating frontal and parietal ROIs, significant differences between stimulation sites (frontal and parietal) were appeared in terms of EF strength in the informed montage (Mean ± SD in V/m: F3 = 0.16 ± 0.07, CP5 = 0.3 ± 0.1). It means that with identical stimulation doses in the frontal and parietal parts of the brain, EF intensity in the parietal is significantly higher than EF intensity in the frontal. However, no significant differences were found in the classics montage between frontal and parietal targets in the left hemisphere (Mean ± SD in V/m: F3 = 0.16 ± 0.07, P3 = 0.18 ± 0.07). Our simulations were replicated for the informed montage to induce balanced EF strength in each stimulation site. Our results showed that using 2 mA in F3 and 0.5 mA in each cathode in the frontal site and 1.2 mA current intensity in CP5 and 0.3 mA in each cathode in the parietal site induce balanced EF strength in both frontoparietal targets. Simulation results for the two last montages in the right hemisphere can be found in Fig.S1.

**Figure 9:**
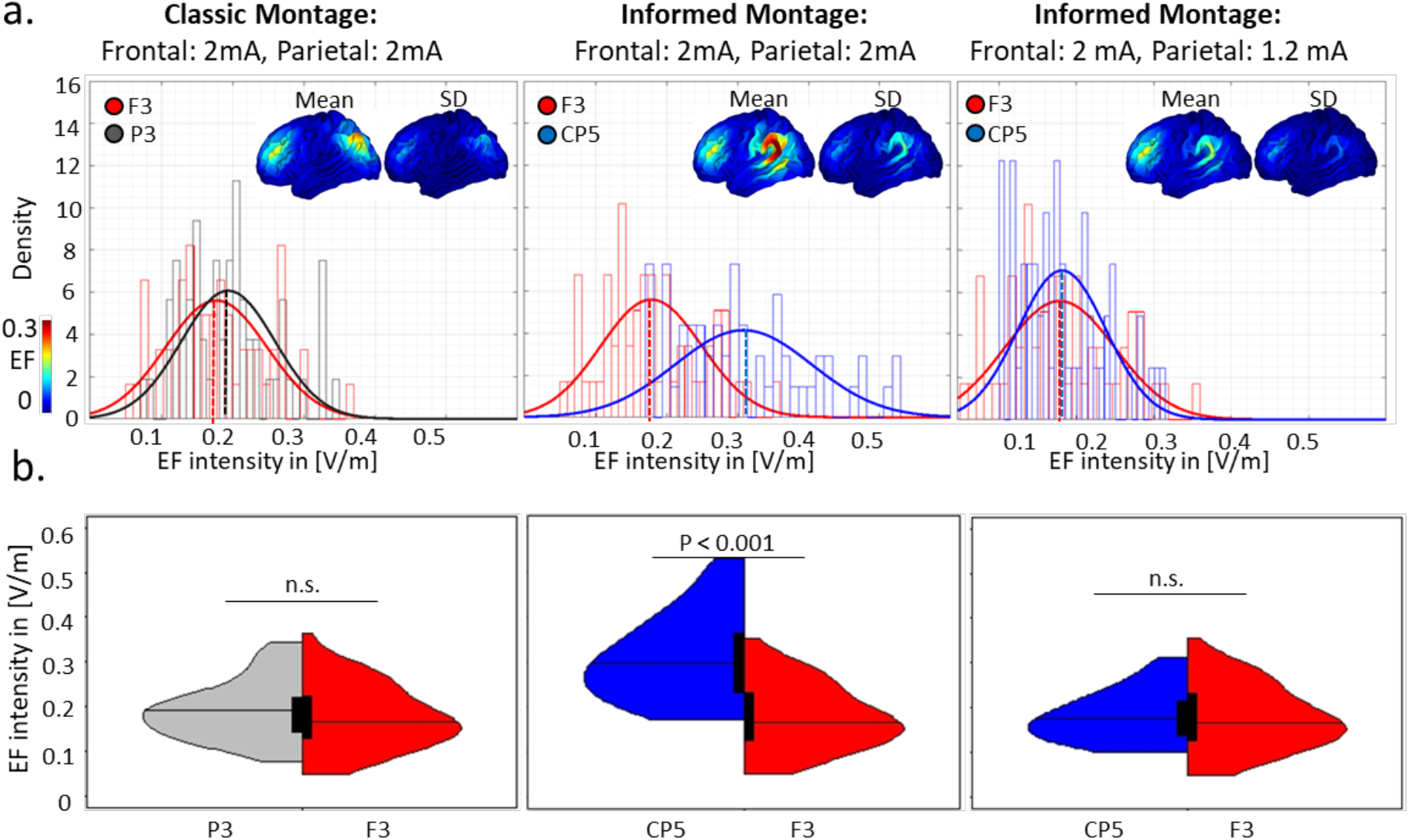
Distribution of the EF in a 10 mm sphere around anode location mapped over the GM (left hemisphere) in the subject and standard space. **a.** Distribution of the EF in volt per meter ([V/m]) are represented for the classic (first column), informed (second column), and informed-modified (third) montages. In the classic and informed montages, the current intensity for anodes was 2 mA and cathodes were 0.5 mA and for informed modified montage in the third column the current intensity for anodes was 2 mA and cathodes were 0.5 mA in the frontal and anode/cathode is 1.2 mA/ 0.3 mA in the parietal site. Anode locations were mapped over the GM for each individual and averaged EFs were extracted from a 10 mm sphere around each anode location in the subject space. The mean and standard deviation of the EFs are also visualized in the fsaverage space based on group-level analysis of the CHMs in standard space. **b.** Split violin plot for EF intensity ([V/m]) as a function of electrode location. Black boxes show the first and third quartile with the median line. Results are calculated for all 60 participants in each 10 mm sphere in stimulation sites in the right hemisphere. Differences between the two montages are shown above the violin plots based on the t-test with FDR correction threshold at P < 0.05. Colors represent electrode locations: F3: in red, P3: in gray, and CP5: in blue. Electrode location for the informed montage: frontal site: anode over F3, cathodes over AF3, F1, F5, FC3. Parietal site: anode over CP5, cathodes over C5, P5, CP3, TP7. Electrode location for the classic montage: frontal site: similar to the informed montage. Parietal site: anode over P3, cathodes over P5, P1, PO3, CP3. <u>Abbreviations</u>: GM: gray matter; SD: standard deviation; CHM: computational head model; n.s.: nonsignificant.

## Discussion

We have described a method for optimizing tES montages with extended targets based on task-based functional connectivity and individualized computational head models. Specifically, this investigation examining electrode locations for modulating frontoparietal network (FPN) with dual-site tES based on simulation of two sets of 4×1 high definition (HD) electrodes in a group of participants with methamphetamine use disorders (MUDs) yielded six main results. First, during the drug cue reactivity task, frontal sites obtained from the head models show psychophysiological interactions with brain regions in the parietal cortex underneath the CP5 and CP6 electrodes. Second, there are significant resting-state functional connectivity and task-based connectivity between F3-CP5 and F4-CP6 (informed montage) considering meth vs. neutral contrast. Third, there was no significant task-based connectivity using brain masks in the classic montage (F3-P3 and F4-P4). Fourth, there was significant task-based and resting-state functional connectivity between frontal seeds and parietal nodes in the ventral attentional network and neighboring (anterior) parietal sites of the FPN based on the Yeo atlas. Finally, interactions with cortico-subcortical brain regions related to the top-down regulation, including the insula, ventral striatum, amygdala, and anterior cingulate cortex, were also found. Taken together, with respect to the functional specificity during tES reported by [24], our results suggest a pipeline for tES electrode montage selection based on fMRI data to modulate cognitive functions in general and drug cue reactivity in specific. Based on our PPI results, F3/F4-CP5/CP6 locations are connected during a cue-reactivity task and might be more susceptible to tES compared to the classic frontoparietal montage (F3/F4-P3/P4). We also recommended current intensity considerations to have balanced EF distribution patterns in targeted stimulation sites.

It has been proposed that tES preferentially modulate a brain area/network that is already activated (e.g., by a specific task) [24]. In this context, target maps for multi-site tES can be defined based on various sources of information such as EEG/MEG or fMRI data [22]. Stimulation targets can be defined based on dynamic interactions between different cortical regions underlie complex brain function [53]. Here, brain regions modulated by a particular task (cue reactivity) were used to determine appropriate electrode locations in FPN modulation instead of using F3/F4-P3/P4 classic montages which are commonly used for modulating FPN functions [16, 33–38, 54]. This approach is in line with previous studies that proposed methods for optimizing stimulation dose based on resting-state functional maps using multi array electrodes that reported potentials for stimulation efficiency improvement compared to non-informed montages [22, 31].

Our ROI-to-ROI gPPI results supported that brain masks in the informed montage have significant interactions between right/left frontal and left/right parietal regions. Between hemispheres involvement (e.g., “left frontal and right parietal” or “right frontal and left parietal”) may facilitate task-driven interactions in the corresponding contralateral cortex via interhemispheric connections that might be related to executive functions [55]. The importance of interhemispheric coupling and its impact on cognitive performance have also been causally tested by applying brain stimulation techniques such as rTMS [56] and tACS [57]. Here, interhemispheric coupling were also investigated in terms of resting-state functional connectivity. The informed montage showed significant positive correlations in interhemispheric connectivity, while negative correlations were found between left frontal and parietal masks (both left and right) in the classic montage. These findings support our seed-to-whole brain gPPI results and suggest that frontal seeds are synchronized with parietal nodes in the informed montage; meanwhile, they are asynchronized (decoupled) in the classic montage. Therefore, it can be hypothesized that informed montage might be more successful in increasing the positive correlation between frontoparietal regions since they are currently working in synchrony with each other during both resting and drug cue reactivity contexts.

The FPN includes a portion of the dorsolateral and medial prefrontal cortex, intraparietal sulcus, precuneus, and cingulate cortex [1]. The ventral attentional network (VAN) or saliency network (SN) includes the lateral prefrontal cortex, anterior cingulate cortex, posterior cingulate cortex, supramarginal gyrus, and parietal operculum. VAN, which is considered in this study, based on Yeo7 atlas parcellation [48] (as well as Schaefer atlas [49]) is named salience ventral attention or salience network (SN), which is related to several salience processes, selective attention behaviors, task switching, and error monitoring [58–60]. Both FPN and VAN have frontal and parietal clusters in their topology. Our exploratory analysis showed that significant frontoparietal connectivity were located next to the parietal nodes in VAN where FPN also had central nodes around this location (supramarginal gyrus). Conversely, parietal nodes in the classic montage were located around the intraparietal sulcus, which was only related to the FPN (and not VAN or SN). Previous neuroimaging studies support that SN and FPN typically show the increase in activation during attention-demanding tasks. However, despite the interaction between SN and FPN, they are eventually distinct and have their own roles [61]. The FPN is a task-positive control network that is linked to external attention and neurocognitive processes, distinct from SN, which is still a task positive network but rather salience-driven. The SN is involved in switching between other large-scale networks, including both task-positive (e.g., FPN) and task-negative (e.g., default mode) networks based on detecting and filtering salient stimuli [1]. Regarding the task-based and resting-state connectivity that we have found in this study, the application of tES to the F3 and F4 sites might be tapping more into the VAN compared to the classic assumption of modulating FPN. The proximity of F3 and F4 sites to the frontal parcels of VAN and connectivity pattern with insula and cingulate sites of the VAN in our results support this consideration.

Based on our data collection sequence (task-based fMRI data immediately after resting-state), resting-state connectivity can be considered as the initial brain-state before encountering drug cues. Correlation analysis between frontoparietal connectivity and VAS alterations during resting-state (negative correlation) and task-based (positive correlation) showed that subjects with lower resting-state frontoparietal connectivity before the cue-reactivity task (as the initial brain-state) experienced higher functional connectivity during meth vs. neutral contrast with a greater level of craving. The negative correlation between resting-state functional connectivity and cue reactivity suggesting that low resting-state connectivity (as the initial brain-state) may decrease the ability to react appropriately to external cues and increase engagement or difficulty to disengagement (or even both) during a drug-cue reactivity task. Our positive correlation between task-based connectivity in VAN (related to the informed montage) and craving score is consistent with previous drug cue reactivity studies that reported craving reflects attentional capture and saliency processing of substance, and participants with a higher level of craving will have difficulty to disengage attention from drug-related cues [62, 63]. Association between the strength of the frontoparietal connectivity and cue reactivity in the informed montage (with parietal nodes next to the VAN) suggesting that the enhanced coupling within the informed montage seeds in the left hemisphere (associated with attentional control) may contribute to cue reactivity in substance users. Therefore, if we hypothesize that the initial strength of the frontoparietal connectivity can affect cue-induced craving (higher connectivity supports lower elicited craving), FPS may help to control drug craving during cue exposure by enhancing connectivity between frontal and parietal parts of the brain and dual-site tACS might be a good candidate for modulating frontoparietal functions. However, future empirical researches are needed to confirm this hypothesis and determine the causal role of FPS and the effectiveness of tES in modulating cue-induced craving.

We found different EF distribution patterns in informed montage compared to the classic electrode arrangement. This might be due to the proximity of the electrodes to the brain and CSF in each montage [64] as well as thickness and composition of the skull in the regions directly underneath the parietal electrodes [64–66]. Furthermore, we found that using the same current intensity across frontal and parietal sites induced significantly different EF distribution patterns in each stimulation site. Therefore, with respect to the non-homologous stimulation sites in FPS, we considered anatomical features of each target site and set the current intensity accordingly. In a recently published paper, the importance of current intensity selection in a frontotemporal tACS protocol was highlighted by considering head modeling approach for determining stimulation dose and different current intensity was used in each stimulation site to have similar EF magnitude in the targeted brain regions [23]. The importance of current intensity selection in multi-channel tACS and the role of CHMs to induce a satisfactory balance between targeted regions in terms of EF strength and focality were also emphasized by Tan et al. [32]. They suggested that in multi-site tACS studies, computational head models should be used for determining stimulation intensities to provide balanced EF distribution patterns in non-homologous targeted regions [32].

Taken together, in this study, a general pipeline was introduced for fMRI informed montage optimization in dual site tES protocols based on using four main steps (Fig.10): (I) determination of the first stimulation site, (II) definition of the fMRI informed activated/connected regions, (III) selection of the second stimulation site, and (IV) optimization of the current intensity in each stimulation site.

**Figure 10:**
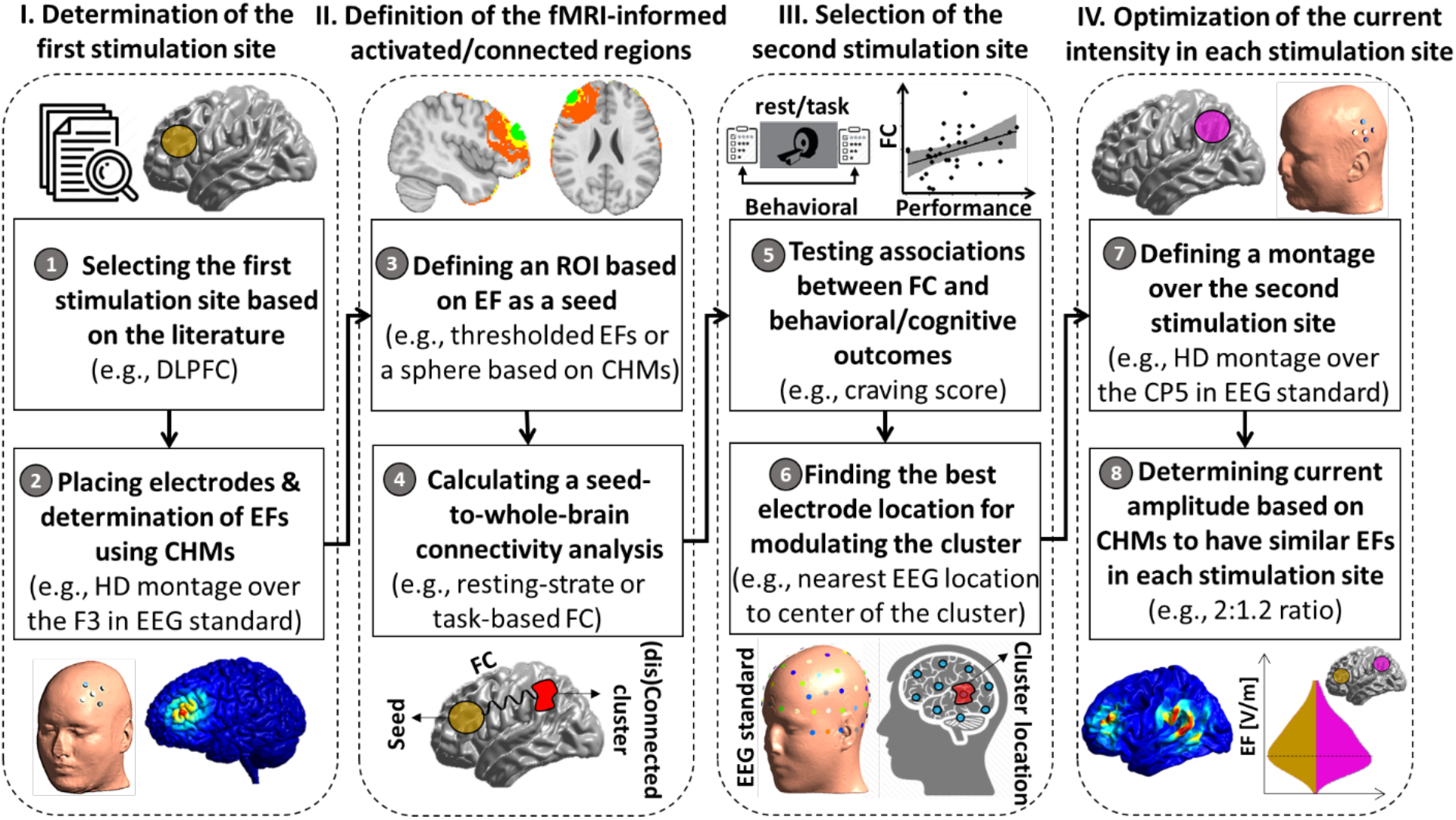
An analytic Pipeline for fMRI Informed Montage Optimization in Dual Site tES. Before applying a dual-site tES for optimally stimulating the target of interest, the main analytical steps are categorized into four main steps. **(I). Determination of the first stimulation site.** (1) The first stimulation site could be determined based on the previous studies for stimulating neural targets of interest (e.g., dorsolateral prefrontal cortex (DLPFC) in frontoparietal stimulation). (2) A set of electrodes will be placed over the neural target (e.g., high definition (HD) electrodes over F3 with a more focal DLPFC targeting compared to the conventional large electrode pads). EF distribution patterns will then be calculated based on computational head modeling (CHM) **(II). Definition of the fMRI-informed activated/connected resions.** (3) A seed region will be defined based on a predefined threshold over the EFs (e.g., a sphere around the maximum EFs). (4) Seed-to-whole-brain connectivity analysis will be performed during rest or task-based fMRI to determine brain regions that are currently activated/connected with respect to the seed region (e.g., changes in functional connectivity during the rest obtained from correlation analysis or task-based (e.g., a cue-reactivity task) connectivity obtained from psychophysiological (PPI) interaction). **(III). Selection of the second stimulation site.** (5) association between behavioral/cognitive outcomes (e.g., craving score before and after a cue-reactivity task) and changes in connectivity (between seed region and currently activated/connected significant cluster) could support validation of the neural target. (6) Based on a predefined criterion (optimality index), the best electrode location for modulating the significant cluster will be determined over the scalp (e.g., EEG 10-10 standard coordinates that are closest to the center of the cluster significantly (dis)connected (from) to the frontal site by drug cues based on Euclidean distance between EEG 10-10 system coordinates and center of the cluster location). **(IV). Determination of the current intensity in each stimulation site:** (7) Electrode montage will be defined based on the area obtained from optimality index (e.g., high definition (HD) electrodes over CP5 as the nearest electrode to the center of the active cluster during a cue-reactivity task). (8) inspired by [32], CHM can be used for determining the ratio of current amplitude at the anodes and cathodes in a way that averaged EF becomes similar in all stimulation sites.

### Limitation and Future Works

Despite positive points, our study has some limitations that could be addressed in future research. We only used two sets of HD eletrodes over frontal and parietal nodes. This includes the need for restriction to a predetermined electrode number. This limitation can be overcome by considering multi-array electrodes (e.g., based on a high-density electrode cap)—which raises the interesting possibility of closed-loop montage optimization using online tES-fMRI or offline tES-EEG trials. Future work is needed to determine the online and offline effects of informed montages for the various types of tES (e.g., tDCS, tACS) on neural/behavioral/clinical outcome measures. It could be investigated whether induced EFs by informed montage link to changes in neural activities. For example, when two main nodes are targeted for frontoparietal synchronization, how other networks and excitatory/inhibitory pathways interact with the targeted brain areas. Furthermore, it would be possible to propose a customized montage for each subject in future work based on individualized brain activity/connectivity at the baseline as suggested in previous TMS studies [67]. Additionally, closed-loop tES-fMRI protocols where ongoing fMRI data is used to optimize stimulation parameters can be used in future studies. Integrating CHMs with the initial brain state obtained from fMRI data can also help design an optimal montage for each person by simultaneously considering the functional and anatomical aspects of stimulation targets. Such considerations have not yet been incorporated into previous dual-site brain stimulation studies.

### Conclusion

The proposed method in this study suggests that our knowledge about activated brain regions during a specific task and connectivity between active brain regions can be used to resolve ambiguity about electrode locations for network-based modulation of the human brain. With the current work, we demonstrated a methodology for montage and dose selection in network-based modulation with tES using task-based fMRI data and modulation of brain connectivity with respect to the cue exposure, adaptable to EEG/MEG connectivity, instead of using typical electrode montages. We specifically considered participants with substance use disorder and drug cue reactivity tasks as an example to effectively activate FPN. However, our pipeline can be applied to other groups of participants, other cognitive tasks, and other large-scale brain networks. More empirical research is needed to support the effectiveness of these measures in clinical applications.

## Author Contribution

G.S., and H.E. designed the study. H.E. and R.K. collected the data. G.S. performed simulations, data analysis, and visualization under H.E. supervision. G.S. wrote the paper with input from R.K., M.P.P, J.B. and H.E. All authors (G.S., R.K., J.B., M.P.P, and H.E.) contributed in manuscript preparation and agreed on the final manuscript before submission.

## Conflict / Declaration of Interest form

The authors declare no competing interest.

## Supplementary Information

### S1. Inclusion and exclusion criteria

The inclusion criteria for this study were: (1) English speaking, (2) diagnosed with MUD (last 12 months), (3) being abstinent from methamphetamine for at least one week, and (4) willing and capable of interacting with the informed consent process. Exclusion criteria included: (1) unwillingness or inability to complete any of the major aspects of the study protocol, including magnetic resonance imaging (i.e. due to claustrophobia), drug cue rating or behavioral assessment, (2) abstinence from methamphetamine for more than 6 months based on self-report, (3) schizophrenia or bipolar disorder based on the MINI [68] interview, (4) active suicidal ideation with intendent or plan determined by self-report or assessment by principal investigator or study staff during the initial screening or any other phase of the study, and (5) positive drug test for amphetamines, opioids, cannabis, alcohol, phencyclidine, or cocaine confirmed by breath analyzer and urine tests.

### S2. fMRI acquisition parameters

Structural and functional MRIs were obtained on one of two identical GE MRI 750 3T scanners at the same site. High-resolution T1 weighted structural images were acquired through magnetization-prepared rapid acquisition with gradient-echo (MPRAGE) sequence using the following parameters: TR/TE = 5/2.012 ms, FOV/slice = 240/0.9 mm, 256×256 matrix, and 186 axial slices, resulting in 0.94×0.94×0.9mm^3^ voxels. High-resolution T2-weighted images were acquired with Fast Spin Echo with radial k-space sampling (PROPELLER) and the following parameters: TR/TE=8108/137.728ms, FOV/slice=240/2mm, 512×512 matrix producing 0.469×0.469×2mm^3^ voxels and 80 coronal slices.

The parameters for task-based and resting-state fMRI data were as follows: the images were acquired by accelerated gradient recalled EPI sequences with Sensitivity Encoding (SENSE) TR/TE = 2000/27 ms, FOV/slice = 240/2.9 mm, SENSE acceleration 2, 96×96 acquisition matrix reconstructed into 1.875×1.875×2.9 mm^3^ voxels, 39 axial slices, and 196 volumes. The parameters for resting-state fMRI data were as follows: the images were acquired by functional imaging EPI sequences with TR/TE = 240/27 ms, FOV/slice = 240/2.9 mm, 128×128 matrix producing 1.875×1.875×2.9 mm voxels, 39 axial slices, and 240 repetitions.

### S3. Individualized computational head models

Individualized CHMs were generated for all participants. SimNIBS 3.2 software pipeline was used for creating CHMs based on FEM. Automated tissue segmentation was performed in SPM 12. The head volume was assigned to six major head tissues (white matter (WM), gray matter (GM), cerebrospinal fluid (CSF), skull, scalp, and eyeballs). The assigned isotropic conductivity values were WM = 0.126 Siemens/meter (S/m), GM = 0.275 S/m, CSF = 1.654 S/m, skull = 0.01 S/m, skin = 0.465 S/m, and eyeballs = 0.5 S/m. The results were visualized using Gmsh and MATLAB 2019b.

In order to compare our suggested electrode configuration with commonly used FPS protocol, a total of 4 dual-site tES electrode montages were simulated and frontoparietal regions in the right and left hemispheres were considered as targeted brain areas. 4×1 HD electrode configuration was considered in each site and, based on using standard EEG cap for each individual in the SimNIBS software, anode electrodes were placed over (1) F3 and P3, (2) F3 and CP5, (3) F4 and P4, and (4) F4 and CP6; with cathode electrodes arranged circularly around each anode. Current densities were corresponding to 2 mA for anodes and 0.5 mA for cathodes. . Electrode location for the informed montage: frontal site: anode over F3, cathodes over AF3, F1, F5, FC3. Parietal site: anode over CP5, cathodes over C5, P5, CP3, TP7. Electrode location for the classic montage: frontal site: similar to the informed montage. Parietal site: anode over P3, cathodes over P5, P1, PO3, CP3. Electrode location for the informed montage: frontal site: anode over F4, cathodes over AF4, F2, F6, FC4. Parietal site: anode over CP6, cathode over C6, P6, CP4, TP8. Electrode location for the classic montage: frontal site: similar to the informed montage. Parietal site: anode over P4, cathodes over P6, P2, PO4, CP4.

### S4. Subregions that were extracted for investigating top-down regulation

These brain regions were extracted based on brainnetome atlas to investigate top-down regulation in response to a drug cue reactivity task.

**Table S1.**
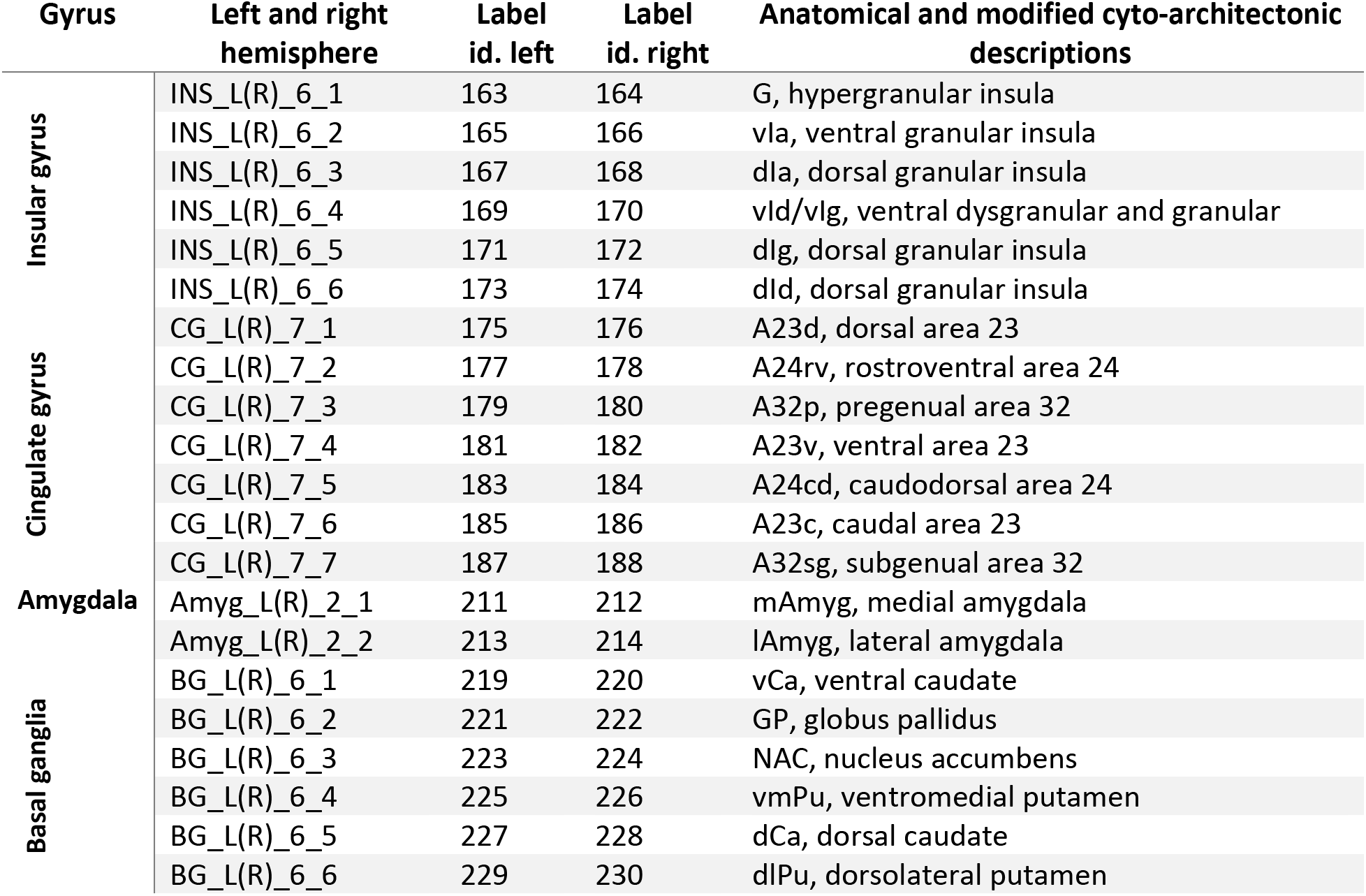
Subregions that were extracted for investigating top-down regulation based on Brainnetome

## Supplementary Figures

**Figure S1:**
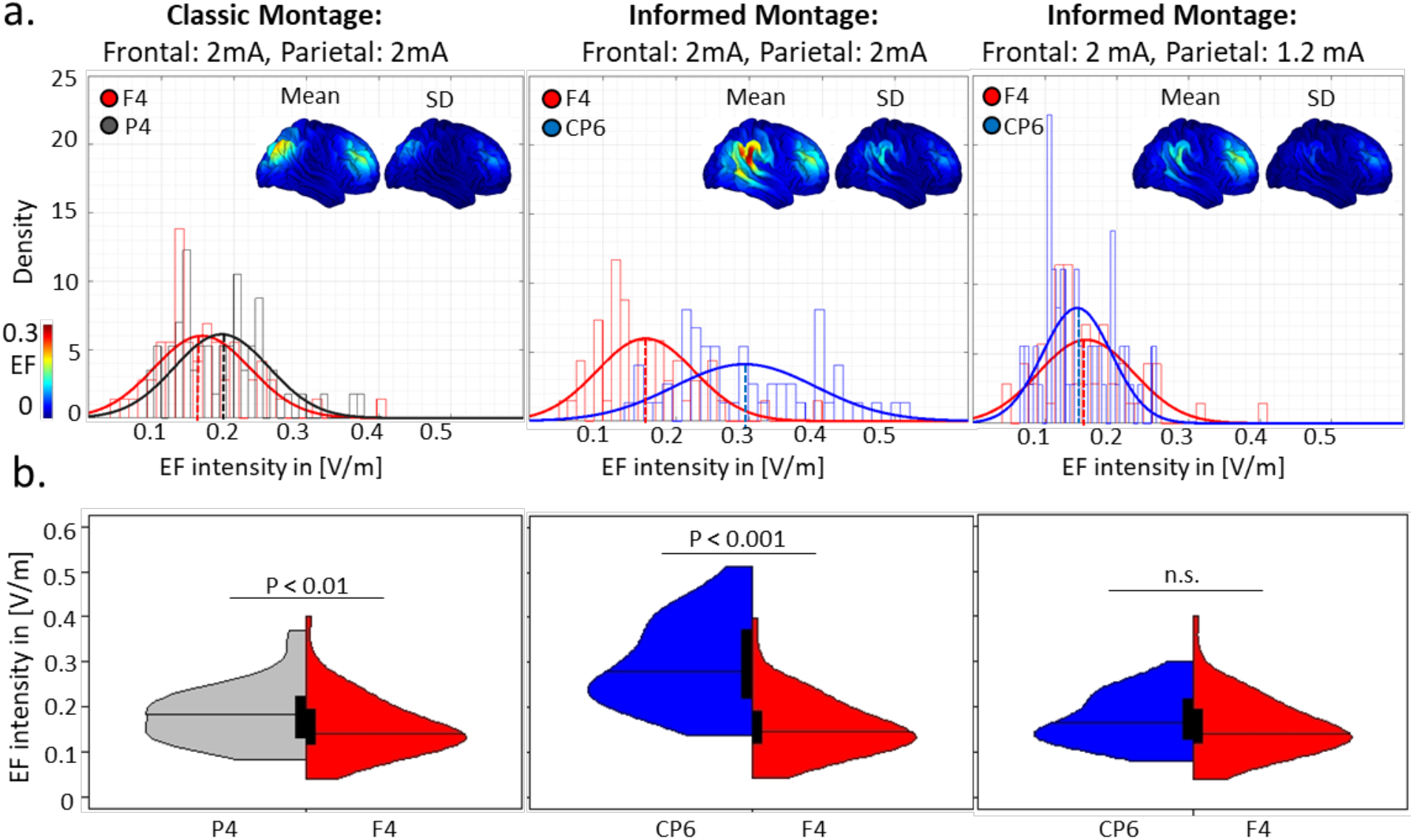
Distribution of the EF in a 10 mm sphere around anode location mapped over the GM (right hemisphere) in the subject and standard space. **a.** Distribution of the EF in volt per meter ([V/m]) are represented for the classic (first column), informed (second column), and informed-modified (third) montages. In the classic and informed montages, the current intensity for anodes is 2 mA and cathodes are 0.5 mA and for informed modified montage in the third column the current intensity for anodes is 2 mA and cathodes are 0.5 mA in the frontal and anode/cathode is 1.2 mA/ 0.3 mA in the parietal site. Anode locations were mapped over the GM for each individual and averaged EFs were extracted from a 10 mm sphere around each anode location in the subject space. The mean and standard deviation of the EFs are also visualized in the fsaverage space based on group-level analysis of the CHMs in standard space. **b.** Split violin plot for EF intensity ([V/m]) as a function of electrode location. Black boxes show the first and third quartile with the median line. Results are calculated for all 60 participants in each 10 mm sphere in stimulation sites in the right hemisphere. Differences between the two montages are shown above the violin plots based on the t-test with FDR correction threshold at P < 0.05. Colors represent electrode locations: F4: in red, P4: in gray, and CP6: in blue. Electrode location for the informed montage: frontal site: anode over F4, cathodes over AF4, F2, F6, FC4. Parietal site: anode over CP6, cathode over C6, P6, CP4, TP8. Electrode location for the classic montage: frontal site: similar to the informed montage. Parietal site: anode over P4, cathodes over P6, P2, PO4, CP4. <u>Abbreviations</u>: GM: gray matter; SD: standard deviation; CHM: computational head model; n.s.: nonsignificant.

## References

1. Marek, S. and N.U. Dosenbach, The frontoparietal network: function, electrophysiology, and importance of individual precision mapping. Dialogues in clinical neuroscience, 2018. 20(2): p. 133.

2. Sheffield, J.M., et al., Fronto-parietal and cingulo-opercular network integrity and cognition in health and schizophrenia. Neuropsychologia, 2015. 73: p. 82–93.

3. Hearne, L.J., J.B. Mattingley, and L. Cocchi, Functional brain networks related to individual differences in human intelligence at rest. Scientific reports, 2016. 6: p. 32328.

4. Cole, M.W., T. Ito, and T.S. Braver, Lateral prefrontal cortex contributes to fluid intelligence through multinetwork connectivity. Brain connectivity, 2015. 5(8): p. 497–504.

5. Ochsner, K.N., et al., Rethinking feelings: an FMRI study of the cognitive regulation of emotion. Journal of cognitive neuroscience, 2002. 14(8): p. 1215–1229.

6. Corbetta, M. and G.L. Shulman, Control of goal-directed and stimulus-driven attention in the brain. Nature reviews neuroscience, 2002. 3(3): p. 201–215.

7. Noudoost, B., et al., Top-down control of visual attention. Current opinion in neurobiology, 2010. 20(2): p. 183–190.

8. Poppe, A.B., et al., Reduced frontoparietal activity in schizophrenia is linked to a specific deficit in goal maintenance: a multisite functional imaging study. Schizophrenia bulletin, 2016. 42(5): p. 1149–1157.

9. Roiser, J.P., et al., Dysconnectivity in the frontoparietal attention network in schizophrenia. Frontiers in psychiatry, 2013. 4: p. 176.

10. Schultz, D.H., et al., Global connectivity of the fronto-parietal cognitive control network is related to depression symptoms in the general population. Network Neuroscience, 2018. 3(1): p. 107–123.

11. Keller, A.S., et al., Paying attention to attention in depression. Translational psychiatry, 2019. 9(1): p. 1–12.

12. Sylvester, C.M., et al., Functional network dysfunction in anxiety and anxiety disorders. Trends in neurosciences, 2012. 35(9): p. 527–535.

13. Ma, Z., et al., Frontoparietal network abnormalities of gray matter volume and functional connectivity in patients with generalized anxiety disorder. Psychiatry Research: Neuroimaging, 2019. 286: p. 24–30.

14. Gürsel, D.A., et al., Frontoparietal areas link impairments of large-scale intrinsic brain networks with aberrant fronto-striatal interactions in OCD: a meta-analysis of resting-state functional connectivity. Neuroscience & Biobehavioral Reviews, 2018. 87: p. 151–160.

15. Monterosso, J.R., et al., Frontoparietal cortical activity of methamphetamine-dependent and comparison subjects performing a delay discounting task. Human brain mapping, 2007. 28(5): p. 383–393.

16. Violante, I.R., et al., Externally induced frontoparietal synchronization modulates network dynamics and enhances working memory performance. Elife, 2017. 6: p. e22001.

17. Jones, K.T., et al., Frontoparietal neurostimulation modulates working memory training benefits and oscillatory synchronization. Brain research, 2017. 1667: p. 28–40.

18. Takeuchi, N., Y. Terui, and S.-I. Izumi, Oscillatory entrainment of neural activity between inferior frontoparietal cortices alters imitation performance. Neuropsychologia, 2021. 150: p. 107702.

19. Hsueh, J.J., et al., Neurofeedback training of EEG alpha rhythm enhances episodic and working memory. Human brain mapping, 2016. 37(7): p. 2662–2675.

20. Shen, J., et al., Real-time fMRI training-induced changes in regional connectivity mediating verbal working memory behavioral performance. Neuroscience, 2015. 289: p. 144–152.

21. Leitao, J., et al., Concurrent TMS-fMRI reveals interactions between dorsal and ventral attentional systems. Journal of Neuroscience, 2015. 35(32): p. 11445–11457.

22. Ruffini, G., et al., Optimization of multifocal transcranial current stimulation for weighted cortical pattern targeting from realistic modeling of electric fields. Neuroimage, 2014. 89: p. 216–225.

23. Reinhart, R.M. and J.A. Nguyen, Working memory revived in older adults by synchronizing rhythmic brain circuits. Nature neuroscience, 2019. 22(5): p. 820–827.

24. Bikson, M. and A. Rahman, Origins of specificity during tDCS: anatomical, activity-selective, and input-bias mechanisms. Frontiers in human neuroscience, 2013. 7: p. 688.

25. Hsu, T.-Y., C.-H. Juan, and P. Tseng, Individual differences and state-dependent responses in transcranial direct current stimulation. Frontiers in human neuroscience, 2016. 10: p. 643.

26. Li, L.M., et al., Brain state and polarity dependent modulation of brain networks by transcranial direct current stimulation. Human brain mapping, 2019. 40(3): p. 904–915.

27. Luft, C.D.B., I. Zioga, and J. Bhattacharya, Anodal transcranial direct current stimulation (tDCS) boosts dominant brain oscillations. Brain Stimulation: Basic, Translational, and Clinical Research in Neuromodulation, 2018. 11(3): p. 660–662.

28. Reato, D., et al., Transcranial electrical stimulation accelerates human sleep homeostasis. PLoS Comput Biol, 2013. 9(2): p. e1002898.

29. Reato, D., et al., Low-intensity electrical stimulation affects network dynamics by modulating population rate and spike timing. Journal of Neuroscience, 2010. 30(45): p. 15067–15079.

30. Stagg, C.J. and M.A. Nitsche, Physiological basis of transcranial direct current stimulation. The Neuroscientist, 2011. 17(1): p. 37–53.

31. Fischer, D.B., et al., Multifocal tDCS targeting the resting state motor network increases cortical excitability beyond traditional tDCS targeting unilateral motor cortex. Neuroimage, 2017. 157: p. 34–44.

32. Tan, J., et al., The importance of model-driven approaches to set stimulation intensity for multi-channel transcranial alternating current stimulation (tACS). Brain Stimulation: Basic, Translational, and Clinical Research in Neuromodulation, 2020. 13(4): p. 1002–1004.

33. Jaušovec, N., K. Jaušovec, and A. Pahor, The influence of theta transcranial alternating current stimulation (tACS) on working memory storage and processing functions. Acta psychologica, 2014. 146: p. 1–6.

34. van Schouwenburg, M.R., T.P. Zanto, and A. Gazzaley, Spatial attention and the effects of frontoparietal alpha band stimulation. Frontiers in human neuroscience, 2017. 10: p. 658.

35. Kleinert, M.-L., C. Szymanski, and V. Müller, Frequency-unspecific effects of θ-tACS related to a visuospatial working memory task. Frontiers in human neuroscience, 2017. 11: p. 367.

36. Pahor, A. and N. Jaušovec, The effects of theta and gamma tACS on working memory and electrophysiology. Frontiers in human neuroscience, 2018. 11: p. 651.

37. van Schouwenburg, M.R., et al., No differential effects of two different alpha-band electrical stimulation protocols over fronto-parietal regions on spatial attention. Frontiers in neuroscience, 2018. 12: p. 433.

38. Jones, K.T., H. Arciniega, and M.E. Berryhill, Replacing tDCS with theta tACS provides selective, but not general WM benefits. Brain research, 2019. 1720: p. 146324.

39. Ekhtiari, H., et al., Methamphetamine and Opioid Cue Database (MOCD): Development and Validation. Drug and Alcohol Dependence, 2020: p. 107941.

40. Dedoncker, J., et al., A systematic review and meta-analysis of the effects of transcranial direct current stimulation (tDCS) over the dorsolateral prefrontal cortex in healthy and neuropsychiatric samples: influence of stimulation parameters. Brain stimulation, 2016. 9(4): p. 501–517.

41. Imburgio, M.J. and J.M. Orr, Effects of prefrontal tDCS on executive function: Methodological considerations revealed by meta-analysis. Neuropsychologia, 2018. 117: p. 156–166.

42. Guo, H., et al., High-definition transcranial direct current stimulation (HD-tDCS) of left dorsolateral prefrontal cortex affects performance in Balloon Analogue Risk Task (BART). Brain and behavior, 2018. 8(2): p. e00884.

43. Nikolin, S., et al., Effects of high-definition transcranial direct current stimulation (HD-tDCS) of the intraparietal sulcus and dorsolateral prefrontal cortex on working memory and divided attention. Frontiers in integrative neuroscience, 2019. 12: p. 64.

44. Thielscher, A., A. Antunes, and G.B. Saturnino. Field modeling for transcranial magnetic stimulation: a useful tool to understand the physiological effects of TMS? in 2015 37th annual international conference of the IEEE engineering in medicine and biology society (EMBC). 2015. IEEE.

45. Nielsen, J.D., et al., Automatic skull segmentation from MR images for realistic volume conductor models of the head: Assessment of the state-of-the-art. Neuroimage, 2018. 174: p. 587–598.

46. Whitfield-Gabrieli, S. and A. Nieto-Castanon, Conn: a functional connectivity toolbox for correlated and anticorrelated brain networks. Brain connectivity, 2012. 2(3): p. 125–141.

47. McLaren, D.G., et al., A generalized form of context-dependent psychophysiological interactions (gPPI): a comparison to standard approaches. Neuroimage, 2012. 61(4): p. 1277–1286.

48. Yeo, B.T., et al., The organization of the human cerebral cortex estimated by intrinsic functional connectivity. Journal of neurophysiology, 2011.

49. Schaefer, A., et al., Local-global parcellation of the human cerebral cortex from intrinsic functional connectivity MRI. Cerebral cortex, 2018. 28(9): p. 3095–3114.

50. Öner, S., Neural substrates of cognitive emotion regulation: a brief review. Psychiatry and Clinical Psychopharmacology, 2018. 28(1): p. 91–96.

51. Ochsner, K.N., et al., Bottom-up and top-down processes in emotion generation: common and distinct neural mechanisms. Psychological science, 2009. 20(11): p. 1322–1331.

52. Fan, L., et al., The human brainnetome atlas: a new brain atlas based on connectional architecture. Cerebral cortex, 2016. 26(8): p. 3508–3526.

53. Shafi, M.M., et al., Exploration and modulation of brain network interactions with noninvasive brain stimulation in combination with neuroimaging. European Journal of Neuroscience, 2012. 35(6): p. 805–825.

54. Polanía, R., et al., The importance of timing in segregated theta phase-coupling for cognitive performance. Current Biology, 2012. 22(14): p. 1314–1318.

55. Kornfeld, S., et al., Resting-state connectivity and executive functions after pediatric arterial ischemic stroke. NeuroImage: Clinical, 2018. 17: p. 359–367.

56. Riddle, J., et al., Causal evidence for a role of theta and alpha oscillations in the control of working memory. Current Biology, 2020.

57. Helfrich, R.F., et al., Selective modulation of interhemispheric functional connectivity by HD-tACS shapes perception. PLoS Biol, 2014. 12(12): p. e1002031.

58. Menon, V., Large-scale brain networks and psychopathology: a unifying triple network model. Trends in cognitive sciences, 2011. 15(10): p. 483–506.

59. Menon, V. and L.Q. Uddin, Saliency, switching, attention and control: a network model of insula function. Brain Structure and Function, 2010. 214(5-6): p. 655–667.

60. Fox, M.D., et al., Spontaneous neuronal activity distinguishes human dorsal and ventral attention systems. Proceedings of the National Academy of Sciences, 2006. 103(26): p. 10046–10051.

61. Chen, A.C., et al., Causal interactions between fronto-parietal central executive and default-mode networks in humans. Proceedings of the National Academy of Sciences, 2013. 110(49): p. 19944–19949.

62. Heitmann, J., et al., Attentional bias for alcohol cues in visual search—Increased engagement, difficulty to disengage or both? PloS one, 2020. 15(1): p. e0228272.

63. Heitmann, J., N.C. Jonker, and P.J. de Jong, A Promising Candidate to Reliably Index Attentional Bias Toward Alcohol Cues–An Adapted Odd-One-Out Visual Search Task. Frontiers in Psychology, 2021. 12: p. 102.

64. Opitz, A., et al., Determinants of the electric field during transcranial direct current stimulation. Neuroimage, 2015. 109: p. 140–150.

65. Ciechanski, P., et al., Modeling transcranial direct-current stimulation-induced electric fields in children and adults. Frontiers in human neuroscience, 2018. 12: p. 268.

66. Mikkonen, M., et al., Cost of focality in TDCS: Interindividual variability in electric fields. Brain stimulation, 2020. 13(1): p. 117–124.

67. Cash, R.F., et al., Personalized connectivity-guided DLPFC-TMS for depression: Advancing computational feasibility, precision and reproducibility. Human Brain Mapping, 2021.

68. Sheehan, D.V., et al., The Mini-International Neuropsychiatric Interview (MINI): the development and validation of a structured diagnostic psychiatric interview for DSM-IV and ICD-10. The Journal of clinical psychiatry, 1998.

